# Cyclic peptides can engage a single binding pocket through multiple, entirely divergent modes

**DOI:** 10.1101/850321

**Authors:** Karishma Patel, Louise J Walport, James L Walshe, Paul Solomon, Jason K K Low, Daniel H Tran, Kevork S Mouradian, Ana P G Silva, Lorna Wilkinson-White, Jacqueline M Matthews, J Mitchell Guss, Richard J Payne, Toby Passioura, Hiroaki Suga, Joel P Mackay

## Abstract

Cyclic peptide display screening techniques can identify drug leads and biological probes with exceptional affinity and specificity. To date, however, the structural and functional diversity encoded in such peptide libraries remains unexplored. We have used the Random nonstandard Peptide Integrated Discovery (RaPID) system to develop cyclic peptide inhibitors of several acetyllysine-binding bromodomains from the Bromodomain and Extra-Terminal domain (BET) family of epigenetic regulators. These peptides have very high affinities for their targets and exhibit extraordinary selectivity (up to 10^6^-fold), making them the highest-affinity and most specific BET-binding molecules discovered to date. Crystal structures of 13 distinct peptide-bromodomain complexes, which all target the acetyllysine-binding pocket, reveal remarkable diversity in both peptide structure and binding mode, and include both α-helical and β-sheet type structures. The peptides can exhibit a high degree of structural pre-organization and bivalent binding of two BDs by one peptide was common, flagging the potential for a new direction in inhibitor design that could bring stronger discrimination between BET-family paralogues. Our data demonstrate for the first time the enormous potential held in these libraries to provide a wide array of modes against a single target, maximizing the opportunity to attain high potency and specificity ligands to a wide variety of proteins.

## Introduction

Cyclic peptides are emerging as a promising class of ligands for challenging biological targets that are capable of blocking a wide range of interactions, including both protein-protein and protein-nucleic acid interactions^1^. This promise stems, in major part, from their size, chirality and dense functionality, which allows development of molecules with exquisite potency and selectivity. A further benefit of cyclic peptides as ligands for protein targets is that these can be rapidly discovery through library technologies (*e.g.*, phage display and mRNA display)^2^. Indeed, such display platforms have been used to isolate macrocyclic peptide ligands against diverse protein targets, including enzymes, proteases, cellular receptors, and growth factors^3–8^. Almost universally, these *de novo* (*i.e.* not derived from known natural peptides) peptides exhibit significantly greater potency and selectivity for their target than other known ligands. However, while the number of examples of successful *de novo* cyclic peptide screens is growing, our understanding of the molecular basis for these interactions – and therefore our appreciation of the ‘binding space’ that such peptides operate in – is relatively poor^9^. Although a number of co-crystal structures have been reported, there has been no systematic analysis of the structural diversity available in typical peptide libraries that would permit discrimination between closely related paralogous protein targets.

Members of the Bromodomain and ExtraTerminal domain (BET) family of transcriptional coregulators (Fig. 1a) each contain a tandem pair of bromodomains (BDs) that recognise acetylated lysine residues in histones and transcription factors^10, 11^. Family members BRD2, −3 and −4 are expressed in a wide range of tissues where they play important – though only partially defined – roles as transcriptional co-regulators; BRD-T plays a similar role in the testes. Inhibition of the BET family of bromodomains has shown considerable promise for a range of different pathologies^12–16^. Indeed, more than two dozen clinical trials are currently underway against a variety of malignancies including acute myeloid leukemia, lymphoma and ER+ breast cancer^17–19^. Despite heavy academic and industry investment in this area, however, the generation of paralogue-selective inhibitors has proved elusive because of the extremely high sequence similarity between the bromodomains of family members (90–95% within the acetyllysine binding pocket residues). This lack of selectivity not only raises issues for the use of BET BD inhibitors as therapeutics, but also limits our ability to probe the mechanisms by which individual BET family members direct gene expression^17^.

**Figure 1.**
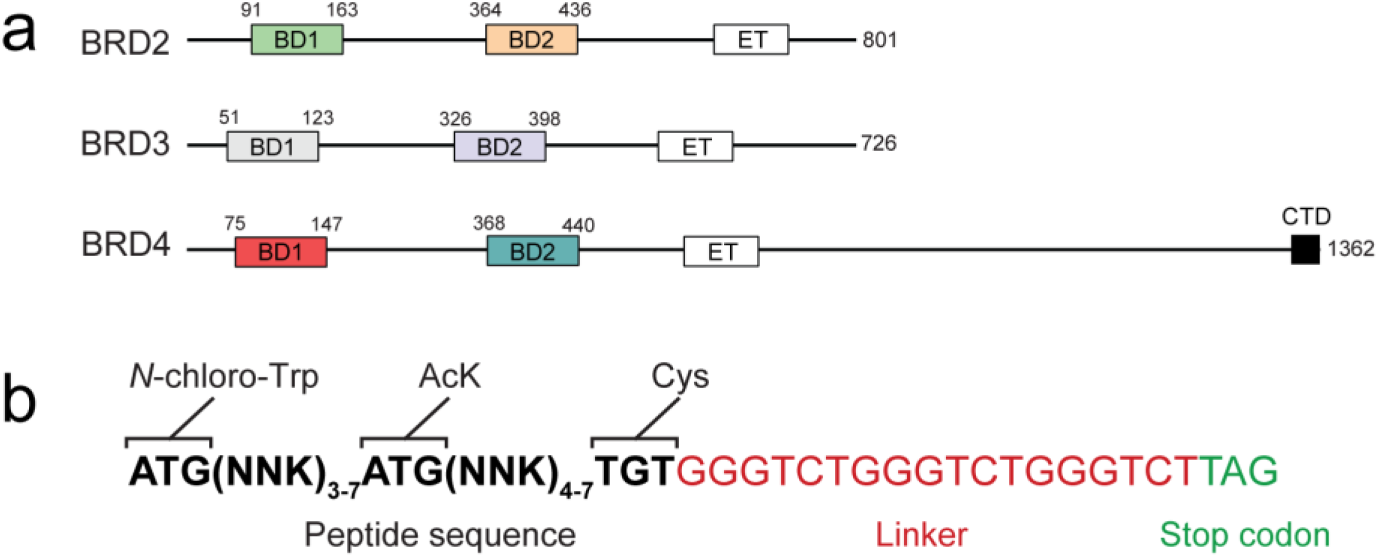
Domain architecture of the BET family and RaPID warhead library design. **a.** Domain topology of human BRD2, −3 and −4. BD1 and BD2 are bromodomains, ET is the extra-terminal domain, and CTD is the C-terminal domain found in BRD4. Residue ranges for the BDs are indicated. **b.** Outline of the warhead library design.

Here we explored the capacity of cyclic peptide library screening to generate high-affinity, selective BET BD ligands. Using a RaPID (Random non-standard Peptides Integrated Discovery) screening approach, which allows the identification of cyclic peptide ligands from extremely large mRNA-encoded libraries (>10^12^ individual members, Supplementary Fig. 1a), we identified numerous very high affinity ligands to three BET family BDs^20^. These ligands, a number of which have sub-nanomolar affinities, exhibited greater selectivity than known inhibitors. Crystal structures of 13 distinct peptide-bromodomain complexes revealed highly diverse structures and binding modes amongst the cyclic peptide ligands. These data are the first demonstration that *de novo* cyclic peptide ligands are capable of binding through multiple, entirely divergent modes to a single pocket.

## Results

### Identification of high affinity cyclic peptide ligands to BRD3-BD1

To identify *de novo* cyclic peptide ligands to BET-family bromodomains, we initially carried out a mRNA display-based RaPID screen against the N-terminal BD of BRD3 (BRD3-BD1). The peptide library for this screen was biased towards substrate-competitive peptides by genetic code reprogramming to replace methionine with acetyllysine (AcK) at the AUG codon^21^. To ensure that at least one AcK was included in each peptide, we designed a “warhead” library with 3–7 degenerate codons flanking each side of a fixed AcK position (Fig. 1b and Supplementary Fig. 1b). The initiating AUG codon was reprogrammed to replace *N*-formylmethionine (fMet) with *N*-chloroacetyl tryptophan to produce macrocyclic peptides through spontaneous reaction with the thiol side chain of a downstream cysteine residue (Fig. 1b and Supplementary Fig. 1b).

The resulting library of >10^12^ cyclic peptides was panned against biotinylated BRD3-BD1 immobilised on streptavidin beads. Following five rounds of selection, the libraries generated from the final three rounds were then sequenced, demonstrating a high degree of sequence enrichment (Supplementary Table 1). As designed, the peptides identified each contained at least one AcK at the fixed central position, but interestingly, many also contained one or more additional AcK residues (Supplementary Table 1). Notably, several biological substrates for BET BDs also bear more than one AcK, with similar spacing to that observed in our selection data^11, 22^. We selected the top two hits as well as the shortest peptide in the top 50 (the 7^th^ hit) for detailed structural and functional studies (Fig. 2a). These peptides were designated **3.1A**, **3.1B** and **3.1C**, and were each synthesised using Fmoc-strategy solid-phase peptide synthesis and cyclized via thioether formation.

**Figure 2.**
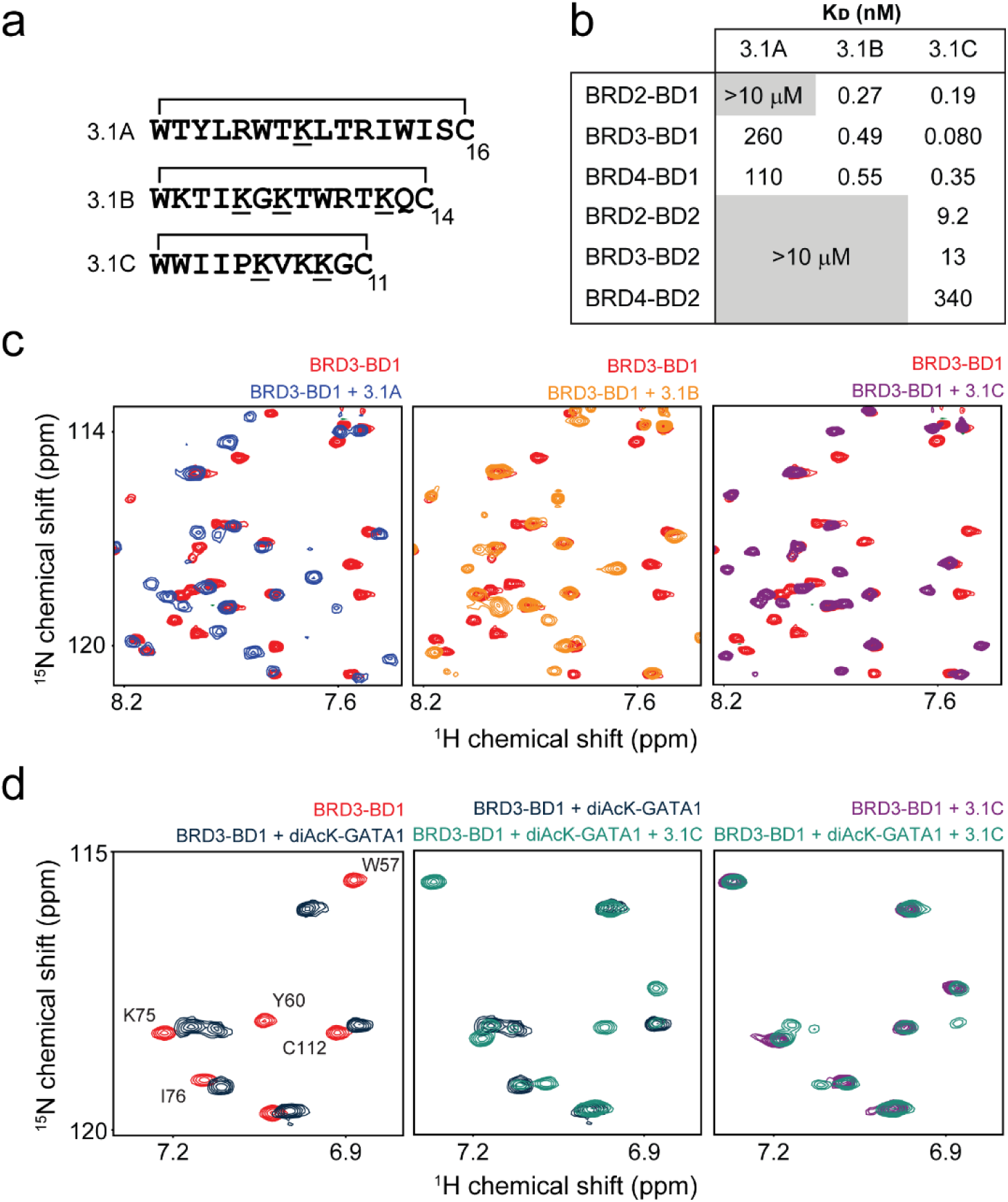
RaPID selection yields cyclic peptides with high affinity and specificity for BRD3-BD1. **a.** Sequences of three peptides selected against BRD3-BD1. Acetylated lysines are underlined and the residues that form the macrocycle are indicated. **b.** Table of K_D_ values for the binding of each BET BD to the peptides **3.1A**, **3.1B** and **3.1C**, measured by SPR. Affinities are given as geometric means of 2– 5 measurements. We estimate the standard deviation of each mean to be ∼40% of the mean. Grey boxed values indicate combinations for which no reproducible signals were observed in SPR experiments at the concentrations used. **c.** Sections of ^15^N-HSQC spectra of BRD3-BD1 either alone (*red*) or in the presence of one molar equivalent of **3.1A** (*left panel, blue*), one molar equivalents of **3.1B** *(middle panel, orange*) or two molar equivalents of **3.1C** (*right panel, purple*). **d.** *Left panel.* Selected region of ^15^N-HSQC spectra of BRD3-BD1 alone (*red*) and in the presence of six molar equivalents of a diacetylated-GATA1 peptide (*dark blue*, GATA1 peptide ranging residues 302-316 and bearing acetylation marks on Lys312 and Lys314). *Middle panel.* Selected region of a ^15^N-HSQC of BRD3-BD1 following the addition of two molar equivalents of **3.1C** to the BRD3-BD1:diacetylated-GATA1 complex formed in the left panel. *Right panel.* Overlay of the final spectrum obtained in the middle panel with the same region of a ^15^N-HSQC of the BRD3-BD1:**3.1C** complex formed in the absence of GATA1.

### RaPID-selected peptides against BRD3-BD1 display extremely high specificity for BD1 domains

We next assessed the ability of each of the three lead peptides to bind BRD3-BD1. Surface plasmon resonance (SPR) studies demonstrated that these cyclic peptides exhibited very high affinity for this domain (Fig. 2b and Supplementary Fig. 2a). To assess selectivity for other BET BDs, we used SPR to determine the binding affinity of each peptide towards the remaining BDs from BRD2, BRD3, and BRD4 (Fig. 2b). All three peptides exhibited marked selectivity for BD1 domains, with observed binding affinities between 25 and >4000 times tighter than those for BD2 domains. Amongst BD1 domains, peptides **3.1B** and **3.1C** displayed little selectivity, with affinities of each peptide for BRD2-BD1 and BRD4-BD1 within a factor of 2–4 of their affinity for BRD3-BD1 – the domain against which they were selected. Peptide **3.1A**, in contrast, demonstrated some selectivity within the BD1 domains as well, displaying similar affinities for BRD3-BD1 and BRD4-BD1, but no observable interaction with BRD2-BD1.

We next used NMR to probe the interactions made by each of these peptides with BD1- and BD2-type BDs. ^15^N-HSQC spectra of BRD3-BD1 alone and in the presence of one molar equivalent of **3.1A**, **3.1B** or **3.1C** showed significant chemical shift perturbations (CSPs) for a sizeable subset of signals (Fig. 2c, Supplementary Fig. 2b-e), consistent with the SPR data. Spectra of the complexes had narrow lineshapes, consistent with the formation of complexes that are well-ordered, stable, and monodisperse. HSQC titrations with other BD-peptide combinations also corroborated the SPR data. In some cases, selective CSPs were observed for combinations that did not show binding in our SPR experiments (*e.g.,* BRD2-BD1 with **3.1A** and BRD2-BD2 with **3.1B**, Supplementary Fig. 2g). The signals for the titration of BRD2-BD2 with **3.1B** were observed to be in fast exchange throughout the titrations, indicating significantly weaker interactions. Fitting the CSP data for the titration of BRD2-BD2 with **3.1B** to a simple 1:1 binding model yielding a binding affinity of ∼300 µM (Supplementary Fig. 2g), which corresponds to a remarkable selectivity of up to 10^6^-fold for the binding of **3.1B** to BD1 domains. Titration of BRD4-BD2 with peptide **3.1A** yielded no significant CSPs (Supplementary Fig. 2h), concordant with the SPR measurements for the same combination. Given the ability of NMR to detect interactions with KD values as weak as ∼1 mM, these data demonstrate that **3.1A** binds BRD3-BD1 and BRD4-BD1 with a selectivity of ∼10,000-fold over BRD4-BD2.

Comparison of the CSPs observed for the BRD3-BD1 complexes (Supplementary Fig. 2c-e) with the crystal structure of the same BD bound to the diacetylated substrate peptide from GATA1 (Supplementary Fig. 2f) indicated that the cyclic peptides all bind to the canonical AcK binding pocket^22^. To confirm this observation, we recorded a ^15^N-HSQC of BRD3-BD1 bound to a diacetylated GATA1 peptide and then further titrated this complex with **3.1C**. Signals that were diagnostic of GATA1 binding (*e.g.*, W57, Y60) shift upon the addition of **3.1C** until they reach, following the addition of one molar equivalent of the cyclic peptide, the same positions observed for the BRD3-BD1:**3.1C** complex (Fig. 2d). These data confirm that the peptides compete directly with natural BRD3-BD1 partners for binding to the AcK pocket.

### Structures of 3.1C bound to BET BDs reveal a highly ordered peptide and the selection of a native-like AcK-BD interaction

We next determined the X-ray crystal structure of BRD3-BD1 bound to **3.1C** (1.9 Å resolution, PDB ID 6U4A, Supplementary Table 2, Fig. 3a-b). The conformation of the BD in the structure closely resembles that reported previously for the same domain bound to the archetypal pan-BD inhibitor JQ1 (PDB ID 3S91; RMSD over ordered Cα backbone = 0.3 Å), as well as that observed in multiple structures of BRD2-BD1 and BRD4-BD1^12, 23, 24^. Strikingly, one of the two AcK residues of the peptide (**3.1C**-AcK6) is inserted into the canonical AcK pocket and displays a sidechain conformation that closely resembles that observed for natural BD-AcK complexes; Fig. 3c shows an overlay of our structure with that of BRD4-BD1 bound to a diacetylated histone H4(1–12) peptide (PDB ID 3UVW), a natural BET BD substrate, for comparison^25^. The previously recognized ‘shelf’ adjacent to the AcK binding pocket – which is occupied by a second AcK in the BRD4-BD1:histone H4 structure – is occupied by the sidechain of Trp2 in our structure, and this sidechain, together with the N-acetyl moiety of Trp1, sandwiches Trp57 of BRD3-BD1 (Supplementary Fig. 3a)^26^. Instead of occupying this shelf, our second AcK residue, AcK9, extends into an adjacent groove (Supplementary Fig. 3b).

**Figure 3.**
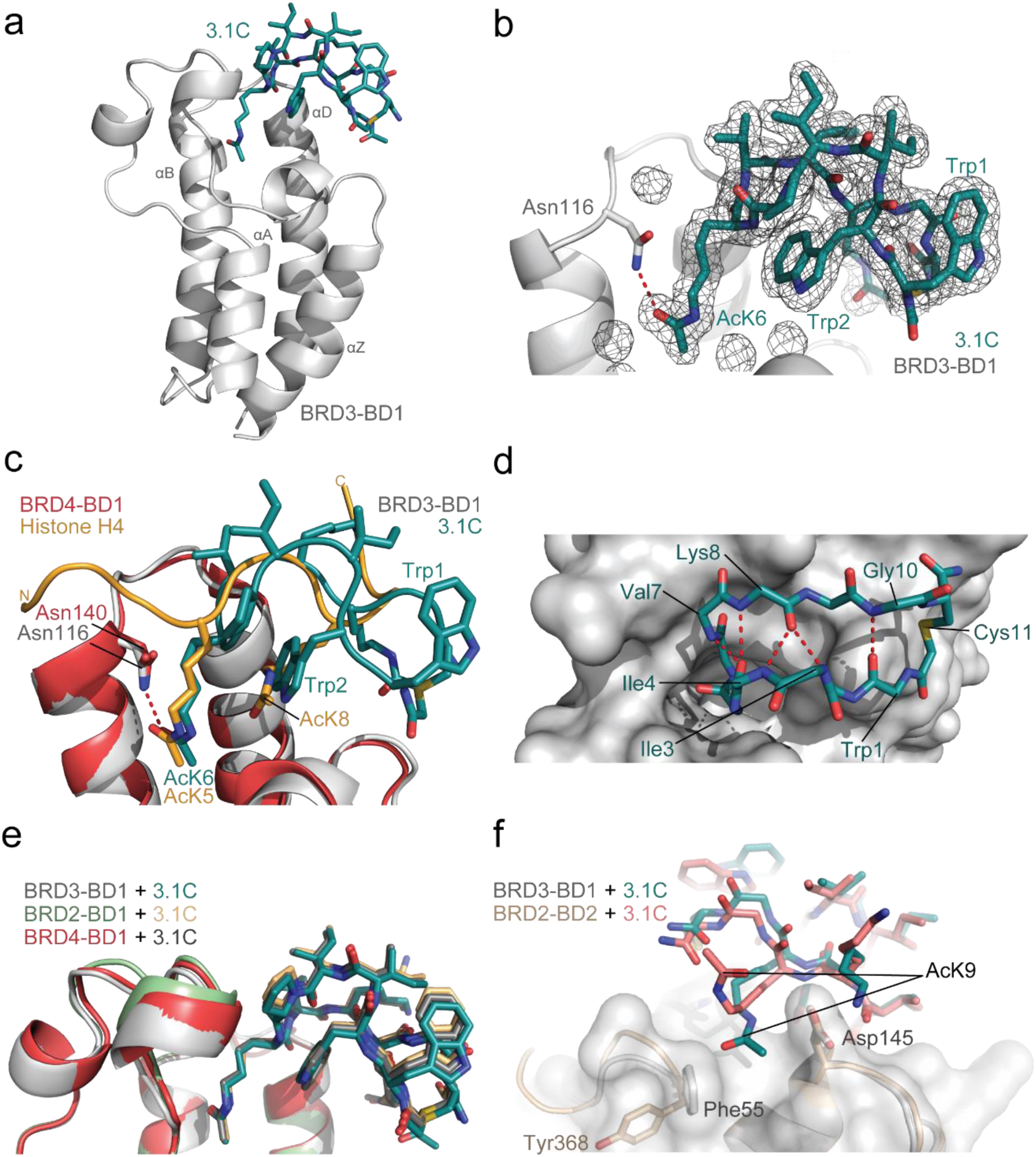
3.1C forms an extended beta structure that inserts an AcK into the canonical binding pocket of BRD3-BD1. **a.** Ribbon representation of the X-ray crystal structure of the BRD3-BD1:**3.1C** complex (1.9 Å resolution, PDB ID 6U4A). The peptide is shown in *teal* and the BD is shown in *grey*. **b.** F_o_–Fc map showing the electron density for **3.1C**, with BRD3-BD1 shown as a cartoon representation. **c.** Overlay of the structure of BRD3-BD1 (*grey*) bound to **3.1C** (*teal*) and of BRD4-BD1 (*coral red*) bound to a diacetylated histone H4(1–12) (*yellow*) peptide. The sidechain conformation of the AcK residues in the binding pocket is very similar. **d.** Internal hydrogen bonding in the backbone of **3.1C**. Hydrogen bonds are indicated as *red* dashed lines. **e.** Overlay of the X-ray crystal structures of BRD3-BD1 (*grey, teal*), BRD2-BD1 (*pale green, light orange*, PDB 2.3 Å resolution, PDB ID 6U61) and BRD4-BD1 (*coral red, dark grey,* 1.7 Å resolution, PDB IDs 6U6K), each in complex with **3.1C**. Structures are overlaid over the heavy atoms of the BDs. **f.** Structure of **3.1C** bound to BRD2-BD2 (*wheat, salmon*,1.5 Å resolution, PDB ID 6U71) overlaid with the structure of **3.1C** bound to BRD3-BD1 (*grey, teal*).

Five intermolecular hydrogen bonds are formed in the complex (including the BRD3-BD1 Asn116 and **3.1C** AcK6 bond characteristic of bromodomain-AcK interactions – Fig. 2b) and 1100 Å^2^ of surface area (across both molecules) is buried. This is low compared to known protein-protein complexes with the same affinity (1500–2500 Å^2^ is typical), pointing to the high binding efficiency that is accessible from cyclic peptide libraries of this type^27^. Another notable feature of the structure is the high degree of internal hydrogen bonding observed for the peptide. The carbonyl group of Ile4 forms a bifurcated hydrogen bond with the amide protons of Val7 and Lys8, the Lys8 carbonyl is similarly hydrogen bonded to the amide protons of Ile3 and Ile4 and the carbonyl of Trp1 forms a hydrogen bond with the amide of Gly10 (Fig. 3d).

We were also able to determine the structure of **3.1C** bound to BRD2-BD1 (2.3 Å resolution, PDB ID 6U61, Supplementary Table 2, Supplementary Fig. 3c) and bound to BRD4-BD1 (1.7 Å resolution, PDB IDs 6U6K, Supplementary Table 3, Supplementary Fig. 3d). The structures overlay very closely with the BRD3-BD1:**3.1C** structure, with the peptide taking up an essentially identical conformation (Fig. 3e). All of the residues in all three structures that contact **3.1C** are completely conserved, consistent with the very similar affinities observed by SPR.

For comparison with the BD1-bound structures, we determined the crystal structure of **3.1C** bound to BRD2-BD2 (1.5 Å resolution, PDB ID 6U71, Supplementary Table 3), to examine the structural basis for the selectivity of this peptide for BD1 domains (Fig. 3f and Supplementary Fig. 3e). Overall, the peptide backbone itself adopts the same conformation and all intermolecular contacts in the AcK-binding region are also conserved. The most apparent differences are a 180° flip of the Trp1 sidechain (Supplementary Fig. 3f) and the nature of the contacts made between AcK9 and the C-terminal part of helix αZ and N-terminal part of helix αD in the BD, more distal to the main AcK pocket (a region less conserved between BD1 and BD2 domains). In the structures with BD1 domains, this AcK lies within a channel formed between conserved Phe (in the loop following αZ) and Asp (αD) residues. A tyrosine residue replaces the Phe in BD2 domains and is positioned in the opposite direction to the Asp, leading to a loss of the aforementioned groove. As a result, AcK9 is oriented towards the solvent in our structure of the peptide with BRD2-BD2 (Fig. 3f and Supplementary Fig. 3g). This difference likely contributes to the significant differences in affinity for the two domains, illustrating the power of such peptides to achieve specificity by probing surfaces beyond the canonical binding pocket.

### 3.1B is a β-hairpin that forms a multivalent interaction with BRD4-BD1 through multiple AcK-mediated interactions

We next determined the X-ray crystal structure of BRD4-BD1 bound to **3.1B** (1.9 Å resolution, PDB ID 6U74, Supplementary Table 4, Fig. 4a). This peptide binds in a very different manner to **3.1C**, while sharing one common central feature: the binding of an AcK (AcK12) in the canonical AcK pocket (Fig. 4b). The peptide forms an ordered β-hairpin with a high degree of internal backbone hydrogen bonding (Fig. 4c). It also contains two additional AcK residues and the structure reveals that, remarkably, these two residues engage a second molecule of BRD4-BD1 (BRD4-BD1-B); AcK5 binds in the AcK-binding pocket and AcK7 contacts the surface of the additional BD (Fig. 4d). The angle at which AcK5 enters the binding pocket differs markedly from the canonical conformation observed for AcK12 (more diagonal than vertical, as depicted in Fig. 4e). Consequently, the AcK hydrogen bond to Asn140 that is characteristic of these complexes cannot form in the same way and is instead mediated by a solvent water molecule (Supplementary Fig. 4a). Some contacts are also made between the surfaces of the two BDs (Fig. 4d).

**Figure 4.**
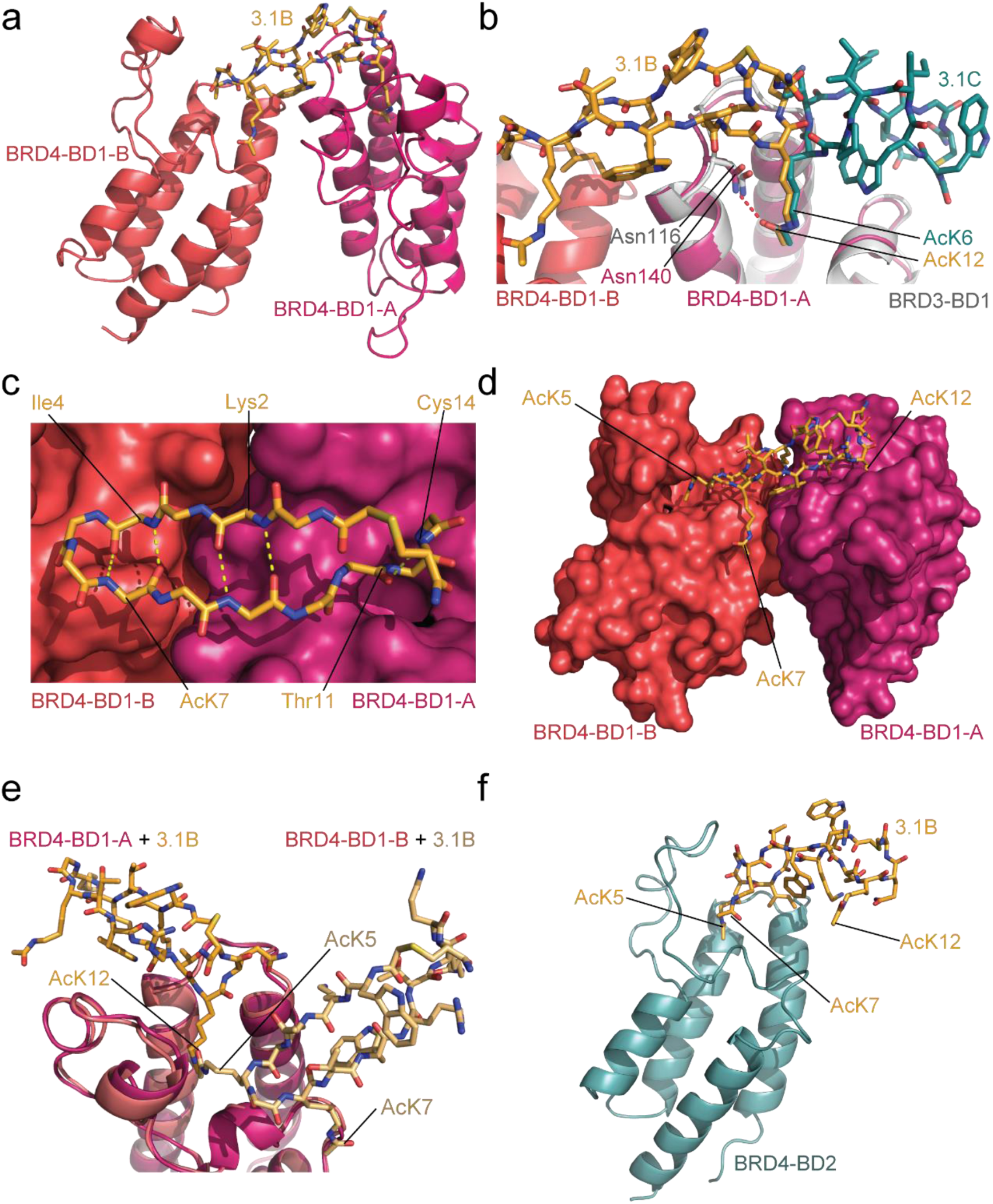
3.1B binds selectively to two BD1 domains through AcK-mediated interactions. **a.** Ribbon diagram of the BRD4-BD1:**3.1B** X-ray crystal structure (1.9 Å resolution, PDB ID 6U74). The peptide is shown in *orange* and the two BDs bound to the peptide are shown in *pink* (BRD4-BD1-A) and *coral red* (BRD4-BD1-B). **b.** Comparison of the binding modes of **3.1B** (*orange*) and **3.1C** (*teal*, bound to BRD3-BD1, *grey*). The two peptides insert an AcK into the AcK-binding pocket in identical conformations but otherwise use completely distinct binding modes. **c.** Internal hydrogen bonding of **3.1B**. Hydrogen bonds are indicated as *yellow* dashed lines. **d.** Positions of the three AcK residues in the **3.1B** complex. AcK5 is partially buried in the AcK binding site of the additional BD (BRD4-BD1-B, *coral red*), whereas AcK7 lies on the surface of that domain. **e.** Comparison of the angle at which AcK12 and AcK5 enter the binding pocket in the two BD subunits in the 2:1 complex. A copy of the BRD4-BD1-B (*coral red*) has been overlaid with BRD4-BD1-A (*pink*) and a copy of **3.1B** (*pale orange*) moved along with it. **f.** Ribbon/stick representation of the X-ray crystal structure of the BRD4-BD2:**3.1B** complex (2.6 Å resolution, PDB ID 6U6L). The peptide is shown in *orange* and the BD is shown in *blue*.

To assess the contributions made by each AcK to BD binding, we synthesized peptides in which each of these three residues was mutated in turn to Ala. Mutation of AcK7 reduced the affinity of **3.1B** for all three BD1 domains by four-fold to ∼2–5 nM (Supplementary Fig. 4b,c), whereas mutation of AcK5 to Ala reduced binding ∼80-fold (KD = 29–35 nM). Mutation of AcK12 to Ala resulted in a complete loss of binding to any BD1 domain under our assay conditions, indicating that this more canonical AcK-pocket interaction provides the largest contribution to binding.

Given this dominant contribution of the AcK12-pocket interaction over the other AcK interactions, we used size exclusion chromatography coupled to multi-angle light scattering (SEC-MALLS) to ask whether equimolar concentrations of the protein and peptide would drive 1:1 complex formation in solution or whether the 2:1 stoichiometry observed in our structure still dominates. Somewhat surprisingly, we established that this complex displays the same 2:1 stoichiometry in solution (Supplementary Fig. 4d), rather than a 1:1 complex (that would have allowed the AcK12 residue to bind in the pocket of each BD). For comparison, we measured the molecular mass of the BRD3-BD1:**3.1C** complex using the same approach (Supplementary Fig. 4d); this complex yielded a molecular mass of 16.6 kDa, in good agreement with the 1:1 stoichiometry observed in our X-ray structure.

Crystal structures of BRD4-BD1 bound to the AcK5→Ala (2.3 Å resolution, PDB ID 6U72, Supplementary Table 4, Supplementary Fig. 4e) or AcK7→Ala peptides (2.6 Å resolution, PDB ID 6U8G, Supplementary Table 5, Supplementary Fig. 4f) were also essentially unchanged from the 2:1 ‘wildtype’ complex. Together, these data suggest that the formation of the ternary complex provides significant additional energetic benefit, underscoring the importance of BD-BD contacts and reminiscent of PROTAC-substrate interactions reported recently^28^.

Finally, we were able to determine the structure of **3.1B** bound to BRD4-BD2, a complex with a much lower affinity than those observed for **3.1B** binding to BD1 domains (2.6 Å resolution, PDB ID 6U6L, Supplementary Table 5, Fig. 4f). In contrast to our observations for peptide **3.1C**, where selectivity between BD1-and BD2-type domains appears to derive from contacts in a region distal to the AcK pocket, the **3.1B** peptide recognizes the BD2 domain in a very different manner. The peptide and BD form a 1:1 complex in which it is AcK5 that is bound in the pocket (*cf*. AcK12 in interactions with BD1 domains). In this case, the interaction surface area is much smaller than observed for the BRD4-BD1 structure, consistent with the substantially lower KD. Furthermore, CSPs observed upon addition of **3.1B** to ^15^N-labelled BRD2-BD2 are consistent with this binding mode (Supplementary Fig. 4g,h), indicating that this peptide binds to all BD2 domains in the same manner. This difference again highlights the ability of RaPID-selected peptides to robustly distinguish BD1-from BD2-type domains.

### Cyclic peptides can also bind selectively to BD2 domains or indiscriminately to all BET BDs with nanomolar affinity

To contrast with our selection against a BD1 domain, we carried out an additional RaPID selection using the same library design but with BRD3-BD2 as the bait. The three most highly enriched peptides (**3.2A**, **3.2B**, and **3.2C**, Fig. 5a) were synthesised, and their BD-binding properties characterised by SPR (Fig. 5b). Peptide **3.2A** proved to be problematic in SPR assays. Surprisingly, strong binding was not observed to BRD3-BD2, against which the peptide was selected in the SPR assay, although it did reproducibly show an interaction with BRD4-BD1 (Fig. 5b). In ^15^N-HSQC experiments, BRD3-BD1 formed a well-defined complex with **3.2A**, with what we estimate to be μM affinity (Supplementary Fig. 5a). However, the addition of the peptide to BRD3-BD2 resulted in the disappearance of most signals, consistent with the formation of soluble aggregates (*e.g.*, Supplementary Fig. 5b) and the poor behaviour of the peptide in SPR experiments.

**Figure 5.**
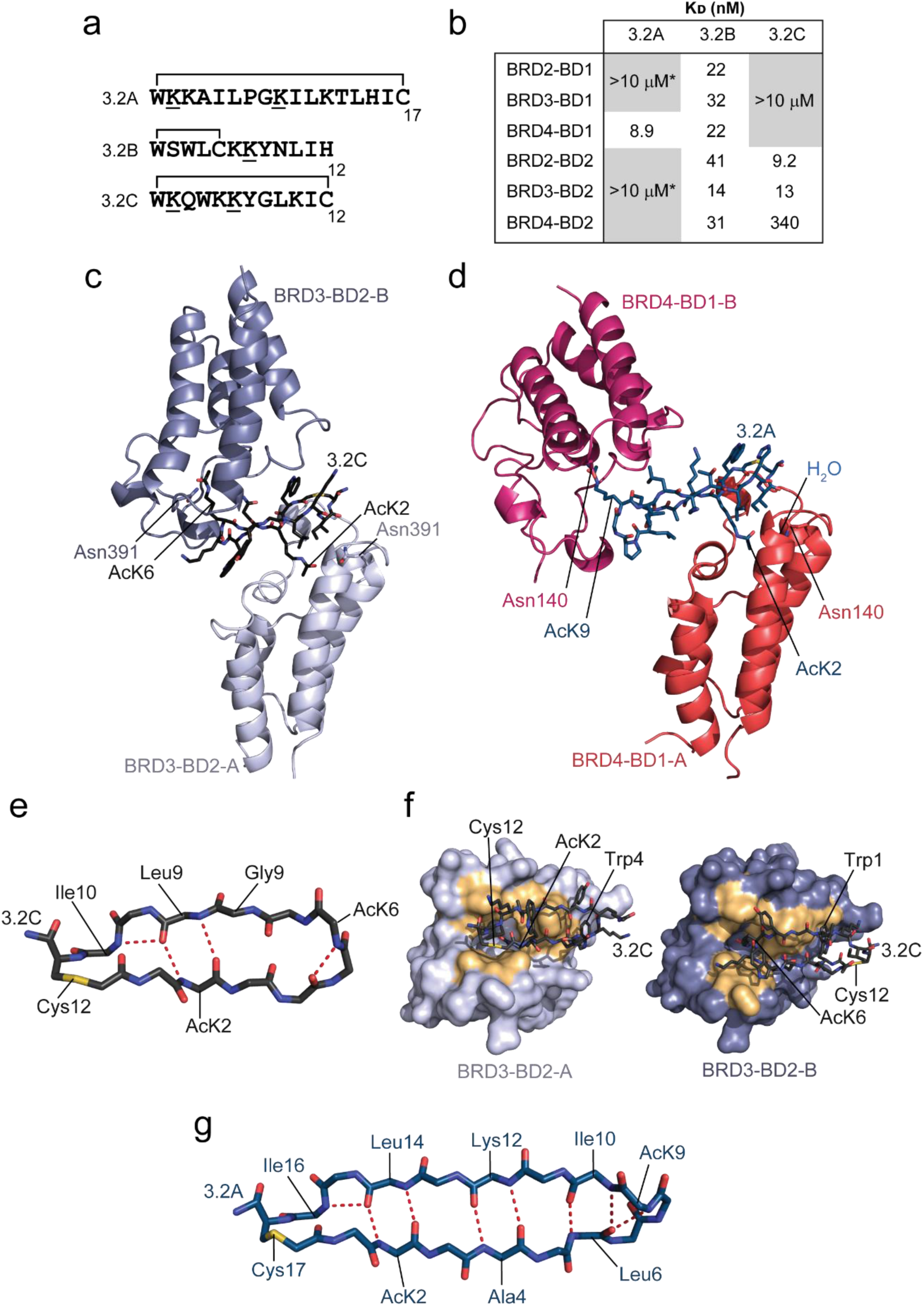
A screen against BRD3-BD2 discovers BD2-selective peptides that bind in a bivalent manner. **a.** Sequences of three peptides selected against BRD3-BD2. Acetylated lysines are underlined and the residues that form the macrocycle are indicated. **b.** Table of K_D_ values for the binding of each BET BD to the peptides **3.2A**, **3.2B** and **3.2C**. Affinities are given as geometric means of 2–5 measurements. We estimate the standard deviation of each mean to be ∼40% of the mean. Grey boxed values indicate combinations for which no reproducible signals were observed in SPR experiments at the concentrations used. **c.** Ribbon diagram of the BRD3-BD2:**3.2C** X-ray crystal structure (2.8 Å resolution, PDB ID 6ULP). The two copies of the BD found in this 2:1 complex are shown in *pale purple* (BRD3-BD2-A) and *dark purple* (BRD3-BD2-B) and the peptide is shown in *charcoal*. AcK residues that enter the binding pocket are labelled. **d.** Ribbon diagram of the BRD4-BD1:**3.2A** X-ray crystal structure (2.0 Å resolution, PDB ID 6U8M). The two copies of the BD found in this 2:1 complex are shown in *coral red* (BRD4-BD1-A) and *pink* (BRD4-BD1-B) and the peptide is shown in *navy*. AcK residues that enter the binding pocket are labelled. **e.** Backbone conformation of **3.2C** with the positions of AcK residues shown and hydrogen bonds indicated as *red* dashed lines. **f.** Comparison of the contact surfaces (*gold*) made by **3.2C** in its interaction with BRD3-BD2-A and BRD3-BD2-B. Similar surfaces mediate both interactions. **g.** Backbone of **3.2A** in the structure with BRD4-BD1. Internal hydrogen bonds are indicated in *red* dashed lines.

Cyclic peptide **3.2B** showed strong binding but relatively little specificity, interacting with each of the six BDs with affinities in the range of ∼10–30 nM (Fig. 5b). In contrast, **3.2C** bound to the BD2 domains with affinities of 10–340 nM but did not bind strongly to any BD1 domain, consistent with selectivities of >10–100-fold between these domains (Fig. 5b).

We were able to obtain X-ray crystal structures of both **3.2C** bound to BRD3-BD2 (2.8 Å resolution, PDB ID 6ULP, Supplementary Table 6, Fig. 5c) and **3.2A** bound to BRD4-BD1 (2.0 Å resolution, PDB ID 6U8M, Supplementary Table 6, Fig. 5d). Like **3.1B**, both peptides bind two copies of the BD. In contrast to **3.1B**, however, almost no direct contacts were observed between the two BDs, suggesting that in these cases crystallization might have promoted the bivalent interaction. Consistent with this idea, SEC-MALLS indicated only weak formation of the 2:1 complex for BRD3-BD2:**3.2C** and exclusively a 1:1 complex for BRD4-BD1 bound to **3.2A** (Supplementary Fig 5c).

In the X-ray structure with BRD3-BD2, **3.2C** adopts an irregular β-hairpin stabilized by several internal backbone amide hydrogen bonds (Fig. 5e). The two AcK residues (AcK2 and AcK6) in the peptide each engage a BD via the AcK pocket, although, in both cases, the N-acetyl moiety lies too far from the conserved Asn (Asn391) to form a direct hydrogen bond. It is likely that water-mediated interactions are formed, but the relatively low resolution of this structure (2.8 Å) prevented us from directly observing solvent water. The two peptide-BD interfaces bury 1120 Å^2^ (BRD3-BD2-A) and 1116 Å^2^ (BRD3-BD2-B) of surface area and the peptide contacts very similar surfaces in the two domains (Fig. 5f) – a surface that significantly overlaps with the contact surface for **3.1C** (Fig. 3d). Despite this overlap in contact surface, the only interaction that is completely conserved is the use of a Trp (Trp4 binding BD-A and Trp1 binding BD-B) to contact Trp322 on the BD.

For **3.2A**, essentially the same face of each of the two BDs is engaged as in the BRD3-BD2:**3.2C** complex, but the angle made between the long axes of the two BDs differs by ∼45°. In the **3.2A** structure, AcK9 forms a direct hydrogen bond with Asn140 in BRD4-BD1-A, whereas the AcK2-Asn140 interaction (with BRD4-BD1-B) is water-mediated (Fig. 5d). A prominent feature of **3.2A** is the high degree of internal backbone hydrogen bonding observed in the structure (Fig. 5g) – nine backbone amide hydrogen bonds form an extremely regular antiparallel β-sheet connected by two turns. Overall, these structures demonstrate that very different sets of amino acids can be used to contact the same protein surface.

### A helical scaffold drives binding of a subset of RaPID-selected peptides to BD2 bromodomains

Cyclic peptide **3.2B** differs from the other library members described above in that it displays a ‘lariat’ topology in which cyclization involves an internal (rather than C-terminal) cysteine and a linear C-terminal tail (Fig. 5a). To assess whether such peptides have different structural features and binding modes, we determined X-ray crystal structures of **3.2B** bound to both BRD2-BD1 (2.1 Å resolution, PDB ID 6U8H, Supplementary Table 7, Fig. 6a) and BRD4-BD2 (2.5 Å resolution, PDB ID 6U8I, Supplementary Table 7, Fig. 6b). Strikingly, **3.2B** forms a single, well-defined α-helix in both structures; this helix contacts the AcK-binding surface of the two BDs with a very similar overall geometry and places its single AcK (AcK7) into the canonical binding pocket. It is notable that the AcK sidechain enters the pocket at a similar diagonal angle to that observed for AcK5 in the peptide **3.1B** binding to the ‘second’ copy of BRD4-BD1 (Fig. 4d, e). Because of this insertion angle, the hydrogen bond made by **3.2B** with the pocket Asn is again mediated by a water molecule (Fig. 6a, b). Residues Trp3, Leu4, Leu10 and Ile11 make hydrophobic contacts with the BDs and Trp3 also contacts the aliphatic part of the AcK7 sidechain, most likely stabilising the conformation of this residue in the binding pocket. Most of these hydrophobic contacts are conserved between the BRD2-BD1 and BRD4-BD2 structures, consistent with the similar affinities observed for the two interactions.

**Figure 6.**
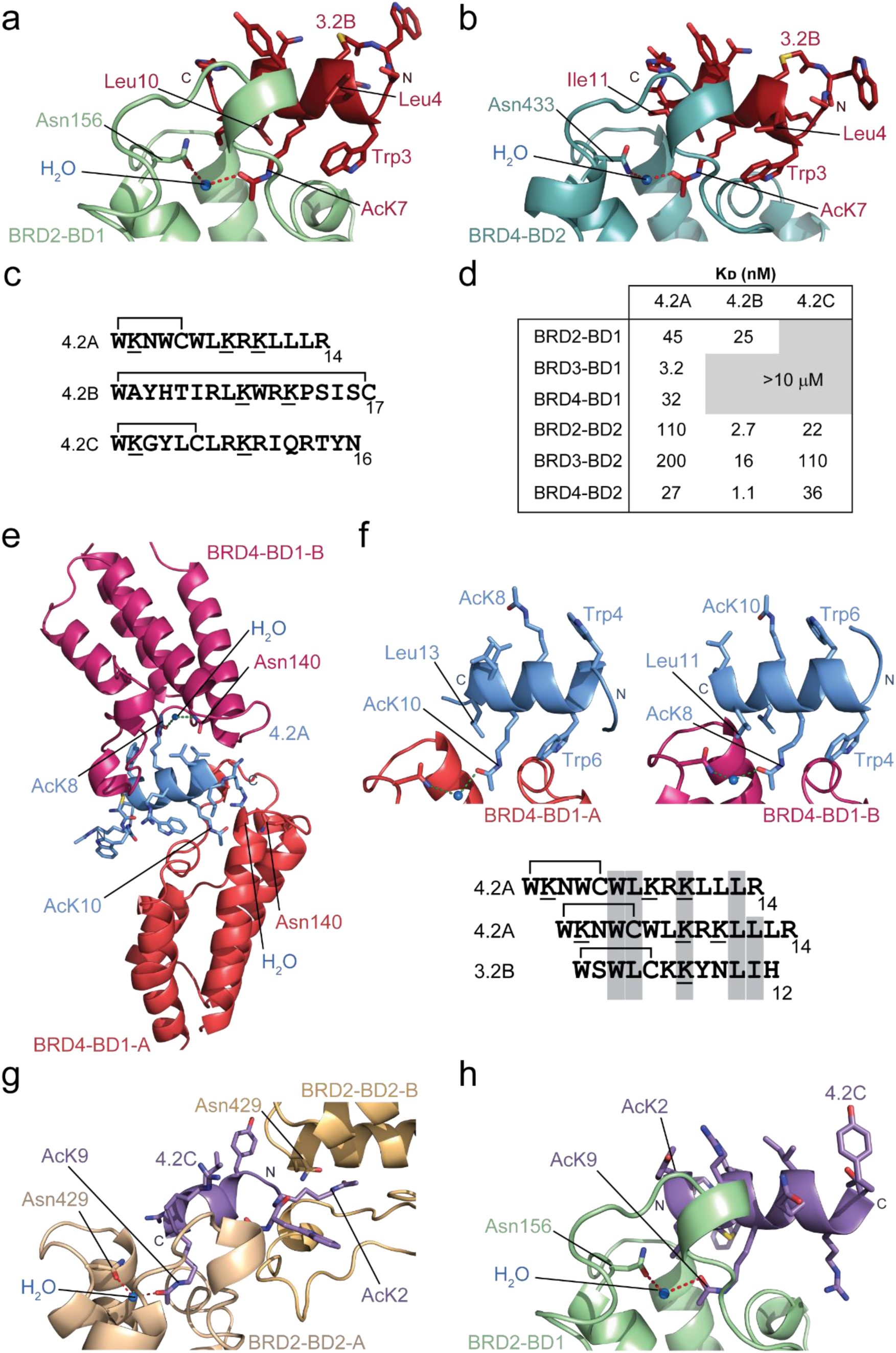
Lariat peptides derived from BD2-directed selections bind in an -helical conformation. **a.** Ribbon diagram of a portion of the BRD2-BD1:**3.2B** X-ray crystal structure (2.1 Å resolution, PDB ID 6U8H). The peptide is shown in *red* and the BD is shown in *pale green*. An ordered water molecule that mediates the interaction between Asn156 of BRD2 and AcK8 of **3.2B** is shown as a *blue sphere* and hydrogen bonding is indicated. Sidechains of **3.2B** are shown as sticks, as is Asn156 from the BD. **b.** Ribbon diagram of a portion of the BRD4-BD2:**3.2B** X-ray crystal structure (2.5 Å resolution, PDB ID 6U8I). The peptide is shown in *red* and the BD is shown in *blue*. An ordered water molecule that mediates the interaction between Asn433 of BRD2 and AcK8 of **3.2B** is shown as a *blue sphere*. Sidechains of **3.2B** are shown as sticks, as is the sidechain of Asn433. **c.** Sequences of three peptides selected against BRD4-BD2. Acetylated lysines are underlined and the residues that form the macrocycle are indicated. **d.** Table of K_D_ values for the binding of each BET BD to the peptides **4.2A**, **4.2B** and **4.2C**. Affinities are given as geometric means of 2–5 measurements. We estimate the standard deviation of each mean to be ∼40% of the mean. Grey boxed values indicate combinations for which no reproducible signals were observed in SPR experiments at the concentrations used. **e.** Ribbon representation of the X-ray crystal structure of BRD4-BD1:**4.2A** (2.2 Å resolution, PDB ID 6ULV). The peptide is shown in *light blue* (with sidechains as sticks) and the BD is shown in *coral red* (BRD4-BD1-A) and *pink* (BRD4-BD1-B). The AcK residues are labelled. **f.** Comparison of mechanism by which **4.2A** interacts with the two BD subunits of the BRD4-BD1:**4.2A** complex. The *left panel* shows the interaction with BRD4-BD1-A. The *right panel* shows the interaction with BRD4-BD1-B. The correspondence between AcK8 and AcK10, L11 and L13, and W4 and W6, can be seen. **g.** Ribbon representation of a portion of the X-ray crystal structure of the BRD2-BD1:**4.2C** complex (2.7 Å resolution, PDB ID 6ULQ). The peptide is shown in *purple* (with sidechains as sticks) and the BD is shown in *pale green*. The direction of the peptide helix is indicated. **h**. Ribbon representation of a portion of the X-ray crystal structure of the BRD2-BD2:**4.2C** complex (2.8 Å resolution, PDB ID 6ULT). The peptide is shown in *purple* (with sidechains as sticks) and the BDs are shown in *wheat* (BRD2-BD2-A) and *pale yellow* (BRD2-BD2-B). The direction of the peptide helix is indicated.

Concurrently, we performed a screen against BRD4-BD2, and measured the binding of the three most enriched sequences (**4.2A**, **4.2B** and **4.2C**), to each of the BDs (Fig. 6c, d). Both **4.2B** and **4.2C** showed significant selectivity for BD2 domains over BD1 domains. **4.2A** bound to BRD4-BD2 with an affinity of 27 nM, ∼4–8-fold tighter than its interaction with other BD2 domains. Surprisingly, affinities of **4.2A** for the three BD1 domains were at least as high as that for BRD4-BD2 – in fact, BRD3-BD1 bound 10-fold tighter (KD = 3.2 nM).

An X-ray crystal structure of the lariat peptide **4.2A** bound to BRD4-BD1 shows that it also forms an α-helix that in this case engages two molecules of BRD4-BD1 in a bivalent complex (2.2 Å resolution, PDB ID 6ULV, Supplementary Table 8, Fig. 6e). SEC-MALLS for this peptide was ambiguous (Supplementary Fig. 5d). SEC-MALLS for **4.2A** binding to BRD4-BD2 indicated a weak 2:1 complex in solution, by contrast experiments for **4.2A** binding to BRD4-BD1 showed only monomer, although the peak was significantly broadened. The peptide interacts with both BD molecules through the insertion of an AcK residue into the binding pocket, and the geometry of both AcKs closely resembles that observed for **3.2B** (featuring a bridging water molecule that mediates the interaction with Asn140; Fig. 6a,b). Remarkably, **4.2A** presents a very similar surface to both BDs in the complex: the peptide sequence harbours two interdigitated Trp-ϕ-X2-AcK-X2-L motifs (ϕ= hydrophobic amino acid) that are directed to opposite sides of the helix to engage the two BDs (Fig. 6f). This interaction surface – along with an additional hydrophobic contact residue observed in one of the **4.2A** interactions – is also conserved in **3.2B** (Fig. 6a,f). The helical Trp-ϕ-X2-AcK-X2-Leu -ϕ motif is therefore a highly favoured BD2-binding unit.

Finally, we solved structures of another lariat type peptide, **4.2C**, bound to a BD1 and a BD2 domain. Fig. 6g shows the structure in complex with BRD2-BD2 (2.8 Å resolution, PDB ID 6ULT, Supplementary Table 9). The peptide is helical and bivalent, recruiting two molecules of BRD2-BD2. Its ‘primary’ interaction has the **4.2C** helix binding in the same location as the other helical peptides but its long axis is ∼30° offset from that of the others. The second interaction is also mediated by an AcK-pocket interaction but with the helix unusually oriented ‘end on’ to the pocket. In contrast, in the structure of **4.2C** bound to BRD2-BD1 (2.7 Å resolution, PDB ID 6ULQ, Supplementary Table 9, Fig. 6h), the peptide takes up the same helical conformation but binds to only a single copy of BRD2-BD1 at the same location as **3.2B** and **4.2A** and with the a similar overall geometry (Fig. 6g). Surprisingly, however, the helix in this case runs in the opposite direction to the other two structures. Together, these structures demonstrate that the class of lariat peptides that we have isolated are predisposed to take up helical conformations and that such peptides can bind BET BDs with affinities that are comparable to those observed for β-hairpin peptides.

### RaPID derived cyclic peptides can be pre-organized for target binding

A notable feature of many of the structures determined here is that the peptides take up very well-ordered α- or β-type conformations with a significant number of intramolecular hydrogen bonds. We asked to what extent these conformations are pre-existing before BD binding by using homonuclear ^1^H NMR spectroscopy to examine several of the peptides in isolation. A 2D NOESY spectrum of **3.1B** showed good dispersion of amide proton resonances and a significant number of medium- and long-range NOEs (Supplementary Fig. 6a), indicating that it is highly ordered in solution. Fig. 7a shows the solution structure of the peptide calculated from these NOE data (PDB ID: 6UXS, Supplementary Table 10). The structure is well defined with the 20 lowest-energy conformers overlaying over heavy backbone atoms with an RMSD of 0.19 Å. Comparison with the structure of **3.1B** in complex with BRD4-BD1 shows that residues Lys2–Thr8 overlay closely whereas Trp1 and Trp9–Cys14 take up a distinct, but well ordered, conformation in the peptide alone. Thus, the peptide is highly ordered in isolation but unexpectedly appears to undergo a significant conformational change – from one well-ordered state to another – upon binding.

**Figure 7.**
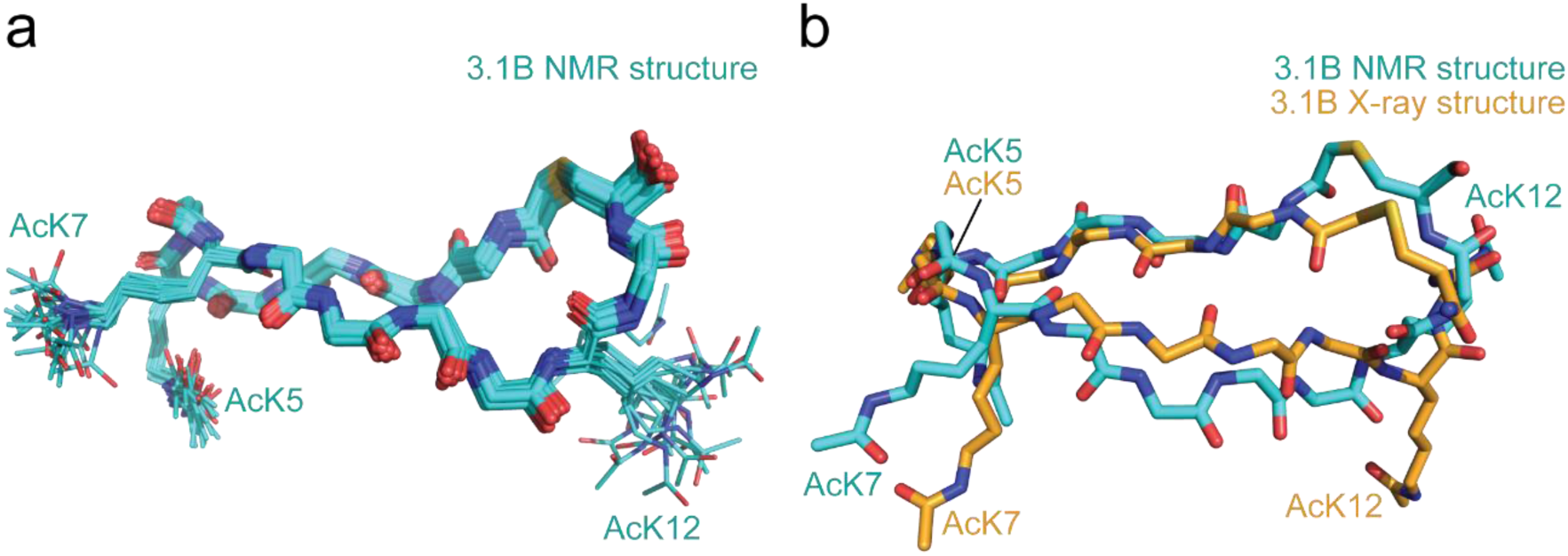
RaPID derived cyclic peptides can be pre-organized for target binding. **a.** Solution structure of **3.1B**. An overlay (over backbone atoms) of the backbones of the 20 lowest energy models is shown, with the sidechains of the three acetyllysine residues shown as sticks. **b.** Alignment (over Cα atoms) of the lowest energy model from the solution structure of **3.1B** with the same peptide from the X-ray crystal structure in complex with BRD4-BD1.

The 1D ^1^H NMR spectrum of **3.2A** and **3.1C** also display well-dispersed amide resonances, consistent with the formation of well-defined and stable structures (Supplementary Fig. 6b). Finally, a NOESY of peptide **4.2C**, which adopts an α-helical structure when bound, displays several amide-amide NOEs, indicating helical propensity in the sequence (Supplementary Fig. 6c). However, the relatively small number of medium- and long-range NOEs suggests that the peptide exhibits significant conformational dynamics rather than a single well-defined conformation. Overall, these data suggest that the library contains numerous peptides that exhibit a significant degree of structural pre-organization, but that peptides with affinities in the nanomolar range are not all highly pre-ordered.

## Discussion

### Cyclic peptide ligands exhibit substantial structural and binding-mode diversity

To the best of our knowledge, the present study is the first comparative investigation of *de novo* cyclic peptide ligands identified from multiple screening campaigns against a series of homologous targets. Our comprehensive structural and biophysical analysis of the most highly enriched peptides from three separate RaPID screens demonstrates that cyclic peptides with a diverse range of structures can bind to even very highly conserved domains with high affinity and specificity. Strikingly, we observed highly diverse structures and binding modes even for cyclic peptide ligands bound to the same pocket of the same BD target.

Despite the high diversity of the library and the high diversity of structures observed in the bound ligands, an AcK residue was found in all cases to bind in the canonical AcK binding pocket, demonstrating both that this is an ‘interaction hotspot’ on the BDs and also flagging the privileged nature of the AcK-pocket interaction with its combination of hydrophobic interactions and a buried amide hydrogen bond. Beyond this commonality, however, the selected peptides adopt a wide range of both alpha and beta type structures that target different features of the BD surface (Supplementary Fig. 7).

It is also notable that the peptides are almost unanimously highly-ordered in the structures of BD-peptide complexes, with only 2–3 residues in a single peptide not observed in the 15 X-ray crystal structures that we describe here. This observation suggests that the highest-affinity peptides for a given target are likely to be those for which much of the peptide takes up a well-defined conformation in the complex and that the interactions are likely to be enthalpy driven. A helical scaffold appears to be particularly favourable for BD2 binding and, strikingly, this property is independent of the direction of the helix, but rather a consequence of the ability of a helix to juxtapose a set of residues to bind the pocket region.

Our structures also reveal that all the peptides themselves adopt compact structures with a high degree of internal hydrogen bonding, features promoted by peptide cyclization. A subset of the peptides also demonstrates a high degree of pre-organisation in the unbound state. This pre-organization is likely a strong contributor to the high affinities that are routinely observed in this and other naïve RaPID screens^29^.

### RaPID-derived cyclic peptides are the highest-affinity and most specific BET-binding molecules discovered to date

BET bromodomain proteins have been therapeutic targets of substantial interest since the discovery that BRD4 is essential for disease maintenance in acute myeloid leukemia nearly 10 years ago^30^. A range of long-running screening and medicinal chemistry campaigns carried out both in industry and in academia have delivered small-molecule BET BD binders with affinities of ∼1–50 nM^31^. The moderate affinities of many of these molecules, combined with their generally low specificity, are potential roadblocks to therapeutic utility and clinical trials to date have had mixed results^31^. Using the RaPID system, we have identified peptides with diverse sequences and structures that bind to BET BDs with affinities of up to 10-fold stronger than the best small molecules. These peptides also display dramatically greater selectivity for BD1s over BD2s (and vice versa) than has been observed previously. The most selective small molecules reported prefer BD1 over BD2 domains by a factor of ∼10–50-fold; in comparison, we describe selectivities of up to ∼10,000-fold or more^31, 32^. Moreover, specificities of up to ∼10-fold were observed in some cases within the BD1 or BD2 subtypes. For example, binding of **3.1A** to BRD2-BD1 could not be observed reproducibly under our SPR conditions, whereas high nanomolar binding was observed for BRD3-BD1 and BRD4-BD1 (Fig. 2b). Similarly, **4.2A** bound BRD3-BD1 30- and 40-fold tighter than BRD4-BD1 and BRD2-BD1, respectively (Fig. 6d). These data suggest that cyclic peptides might ultimately provide a route to achieving the intra-subfamily selectivity that has proved elusive with other classes of small molecules. Further efforts are ongoing to enhance this selectivity.

### Bivalent complexes offer opportunities for more sophisticated BET targeting strategies

In many cases our peptides were found to contain more than one AcK residue. This presence of more than one AcK is a common feature of many bromodomain-binding acetylated transcription factors and histone proteins^11, 22^. Whilst in some cases, as in nature, these multiple AcK residues bind to a single bromodomain, we also observed multiple instances of bivalent bromodomain binding. It is likely that this multi-valency plays a role in the specificity of peptides for particular BDs, especially in cases where there are significant contacts between the two BDs – such as in the interactions between **3.1B** and BD1-type domains.

The observation of bivalent binding also suggests a strategy for the design of peptides (or other molecules) that can simultaneously bind both tandem BDs within a single BET family member (*e.g.*, BRD2-BD1 and BRD2-BD2). There is the potential for such pseudo-ternary complexes to display markedly higher specificity and affinity for one paralogue.

Overall, our findings demonstrate that relatively small macrocyclic peptide ligands can adopt highly diverse conformations during binding to target proteins, and greatly expand the catalogue of *de novo* cyclic peptide-protein crystal structures solved to date. Whereas previous studies have demonstrated that such peptide ligands can adopt recognisable secondary structural features (*e.g.* α-helix or anti-parallel β-sheet) when bound to specific binding pockets, our findings demonstrate for the first time that peptides with diverse structures can bind to the same pocket. This has profound implications for the design of cyclic peptide ligands, since optimal ligands may be structurally unrelated to known ligands in many cases. This finding likely also contributes to the success of RaPID and related strategies for isolating high-affinity ligands.

## Materials and Methods

### Protein expression and purification

The bromodomains from human BRD2 (BD1: 65-194; BD2: 347-455), BRD3 (BD1: 25-147; BD2: 307-419), and BRD4 (BD1: 42-168; BD2: 348-464) were cloned for bacterial expression into the pGEX-6P plasmid for expression as N-terminal GST-fusion proteins and into pQE80L-NAvi for expression as N-terminally His-tagged and biotinylated proteins.

Biotinylated bromodomains were prepared for SPR using the pQE80L-NAvi constructs. Proteins for crystallography, SEC-MALLS, and MST were produced using the pGEX-6P constructs. All bromodomain constructs, except BRD4-BD1, were transformed into competent BL21(DE3) *Escherichia coli* (*E. coli*) cells. LB expression cultures were inoculated with saturated starter cultures (1:100 dilution) grown from single colonies from fresh transformations plates. Expression cultures were incubated at 37 °C, with shaking at 150 rpm, until cultures reach mid-log phase (OD600 of ∼0.6-0.8). At this point, cultures were cooled to room temperature and supplemented with 250 µM IPTG (and 200 µM biotin for pQE80L-NAvi constructs). Expression cultures were transferred to 18 ⁰C for expression for a further ∼20–24 h before harvesting via centrifugation. Biotinylated BRD4-BD1 was expressed using Rosetta2(DE3) *E. coli* cells. Autoinduction expression cultures were inoculated with a starter culture, as described above, and incubated at 37 °C with shaking at 150 rpm. Upon reaching mid-log phase, cultures were cooled to room temperatures and supplemented with 200 µM biotin. Expression cultures were transferred to 18 °C for expression for a further ∼20–24 h before harvesting. All media were supplemented with the appropriate antibiotics. All cultures were harvested via centrifugation at 5000 ×g at 4 ⁰C for 25 min and cell pellets were stored at –20 ⁰C.

^15^N-labelled proteins were expressed by transforming competent BL21(DE3) *E. coli* cells with pGEX-6P constructs. LB cultures were inoculated as described in the previous section. The cultures were harvested and washed in M9 minimal media salts once they reached mid-log phase. The cells were then resuspended in a volume of minimal media, containing ^15^NH4Cl, half the volume of the original LB culture. The cultures were first transferred to 18 ⁰C for 45 min and then expression was induced using 250 µM IPTG. The cultures were incubated at 18 ⁰C for expression for a further ∼20–24 h, with shaking at 150 rpm. All cultures were supplemented with antibiotics, harvested, and stored as described in the previous section.

Pellets from cells expressing proteins for SPR were lysed via sonication in a buffer composed of 50 mM Tris pH 8.0, 500 mM NaCl, 20 mM imidazole, 5 mM β-mercaptoethanol (β-ME), 0.1% Triton X-100, 1× cOmplete EDTA-free protease inhibitor, 10 µg/mL DNase I, 10 µg/mL RNase, and 100 µg/mL lysozyme. The lysate was clarified via centrifugation at 18,000 ×g for 30–60 min. The soluble fraction of the clarified lysate was subjected to immobilised nickel ion affinity chromatography using a 1-mL HisTrap column. The bound protein was eluted using a 20–250 mM imidazole gradient elution. The protein containing eluates from the affinity chromatography step were pooled and concentrated to a small volume. The concentrated sample was subjected to size exclusion chromatography (SEC) using a HiLoad 16/600 Superdex 75 column. Protein was eluted from the column using 50 mM Tris pH 8.0, 150 mM NaCl, and 1 mM dithiothreitol (DTT). Protein containing eluates were pooled and either aliquoted directly or concentrated before aliquoting. Aliquots were snap frozen in liquid nitrogen and stored at –80°C. Protein purification was analysed by SDS-PAGE and monitoring UV absorbance at 280 nm.

Cell pellets for protein for all other purposes were lysed in a buffer composed of 50 mM Tris pH 7.2, 500 mM NaCl, 5 mM β-mercaptoethanol (β-ME), 0.1% Triton X-100, 1× cOmplete EDTA-free protease inhibitor, 10 µg/mL DNase I, 10 µg/mL RNase, and 100 µg/mL lysozyme via sonication. Lysate was clarified via centrifugation at 18,000 ×g for 30–60 min. The soluble fraction of the clarified lysate was subjected to GSH-affinity chromatography using a 5-mL GSTrap column. Bound protein was step eluted with 50 mM Tris pH 7.2, 150 mM NaCl, 10 mM reduced glutathione, and 5 mM β-ME. Protein containing eluates were pooled and incubated with HRV-3C protease overnight at 4 ⁰C to enable cleavage of the GST-tag. The pooled eluate was concentrated and subjected to size exclusion chromatography using a HiLoad 16/600 Superdex 75 column. The protein was eluted from the column in a buffer comprising 10 mM Tris pH 7.2, 100 mM NaCl, and 1 mM dithiothreitol (DTT). Fractions containing the desired protein were pooled and concentrated before aliquoting. ^15^N-labelled protein was concentrated to 5–10 mg/mL and unlabelled protein was concentrated to 10–15 mg/mL. Protein aliquots were snap frozen in liquid nitrogen and stored at –80 ⁰C.

### RaPID screening

DNA sequences containing a T7 polymerase binding site, ribosome binding site, variable length randomised peptide coding region containing a fixed central ‘ATG’ codon for warhead incorporation, (Gly-Ser)3, linker, ‘TAG’ stop codon and sequence for puromycin ligation were constructed by extensions and PCR from oligos purchased from Eurofins Genomics K.K. (Japan) to produce warhead libraries.

Library DNA - TAATACGACTCACTATAGGGTTGAACTTTAAGTAGGAGATATATCCATG(NNK)_m=3_ -7ATG(NNK)_n=4-7_TGTGGGTCTGGGTCTGGGTCTTAGGTAGGTAGGCGGAAA DNA libraries were transcribed to mRNA using T4 RNA polymerase and mRNA libraries were ligated to a puromycin-PEG-DNA splint using T4 RNA ligase following standard reaction conditions. For the first selection round libraries were mixed in the following proportions: (m=3,n=4):(m=4,n=4):(m=4,n=5):(m=5,n=5):(m=5,n=6):(m=6,n=6):(m=6,n=7):(m=7,n=7) = 0.01425:0.45:10:10:7.5:7.5:7.5:7.5

For codon reprogramming, the 3,5-dinitrobenzyl ester of *N^ε^*-acetyl-lysine was synthesised as previously described and aminoacylated onto tRNA^Asn^_CAU_ using dFx (2hr, RT, standard aminoacylation conditions) for incorporation into library peptides^33^. Peptides were initiated with *N*-(chloroacetoxy)-L-tryptophan (ClAc-L-Trp), which was aminoacylated via the cyanomethyl ester onto tRNA^fMet^_CAU_ using eFx (2hr, RT, standard aminoacylation conditions). Flexizymes were prepared as described previously^34^.

RaPID screens were carried out as previously described^35^. Briefly, puromycin-ligated randomised mRNA libraries were *in vitro* translated (30 min, 37 °C then 12 min, 25°C) using a custom transcription/translation mixture containing additional 12.5 µM ClAc-L-Trp-tRNA^fMet^_CAU_ and 25 µM AcK-tRNA^Asn^_CAU_ and lacking methionine and 10-formyl-5,6,7,8-tetrahydrofolic acid^36^. First-round translations were carried out on a 150-µL scale, subsequent rounds on a 5-µL scale. Following addition of 200 mM EDTA, pH 8.0 (15 µL) and reverse transcription with M-MLV RTase, RNase H minus (Promega), blocking buffer was added (50 mM HEPES, 150 mM NaCl, 2 mM DTT, 0.1% Tween-20, 0.2% (w/v) acetylated bovine serum albumin, pH 7.5) and libraries were incubated with magnetic streptavidin bead-immobilised bromodomain (Promega) (200 nM, 30 min). Following washing (156 µL ice-cold 50 mM HEPES, 150 mM NaCl, 2 mM DTT, 0.1% Tween-20, pH 7.5, 3 × 5 min), 400 µL PCR solution was added and retained peptide-mRNA/DNA hybrids were eluted from the beads by heating (95 °C, 5 min). Library enrichment was assessed by quantitative real-time PCR relative to standards and the input DNA library using primers T7g10M_F46 and CGS3-CH.R22. Recovered libraries were amplified using the same primers and used as the input DNA for subsequent selection rounds.

T7g10M_F46 - TAATACGACTCACTATAGGGTTGAACTTTAAGTAGGAGATATATCCCGS3-CH.R22 - TTTCCGCCTACCTACCTAAGAC

Following RaPID selections, double indexed libraries (Nextera XT indices) were prepared from recovered library DNA from rounds 3–5 and sequenced on a MiSeq platform (Illumina) using a v3 chip as single 151 cycle reads^37^. Each DNA sequence was converted to a peptide sequence and ranked by total read number (Supplementary Table 1).

### Peptide synthesis

Peptides were synthesised as C-terminal amides using standard fluorenylmethyloxycarbonyl (Fmoc)-strategy solid-phase chemistry using a Syro I automated synthesiser (Biotage) and NovaPEG Rink Amide resin (Novabiochem). Couplings were performed with HBTU/HOBt (1:1) and 6 equivalents of each amino acid. Double couplings were performed for arginine residues. Following the final amino acid coupling reaction, the Fmoc group was removed and resin incubated with N-(chloroacetoxy)succinimide (0.2 M in DMF, 1 h, RT). Resin was washed with DMF (5 times) and DCM (5 times) and dried *in vacuo*.

Linear peptides were cleaved from the resin by incubation (3 h, RT) with a trifluoroacetic acid (TFA) cleavage cocktail (TFA/2,2’-(ethylenedioxy)diethanethiol/triisopropyl silane/H2O (92.5:2.5:2.5:2.5)) before filtration, concentration *in vacuo* and precipitation with ice-cold diethyl ether. Crude peptides were washed with diethyl ether (5 times), dried and resuspended in DMSO. The pH was raised to >8 using triethylamine to allow cyclisation. Following incubation (1 h, RT), peptides were re-acidified with TFA for purification.

Crude peptides were purified by reverse-phase high-performance liquid chromatography using a Chromolith Prep column (Merck) on a Prominence LC-20AP system (Shimadzu) (Solvent A: 0.1% TFA in H2O, Solvent B: 0.1% TFA in acetonitrile) to >95% purity as determined by mass spectrometry (Supplementary Fig. 8). Purified peptides were reconstituted in DMSO and concentrations determined from their absorbance at 280 nm in 5% DMSO using predicted extinction coefficients.

### Surface plasmon resonance (SPR)

Measurements were conducted on a T200 or S200 (GE Healthcare) and data analysed using the Biacore Insight Evaluation Software. Experiments were performed at 4 °C in single cycle kinetics mode. Biotinylated BET bromodomains were immobilised on a CAP chip (GE Healthcare) with a target density of ∼1000-1500 RU. 50 mM HEPES, 150 mM NaCl, 0.05% Tween-20, 0.1% DMSO, pH 7.5 was used as the running buffer. Between cycles the chip was regenerated following the manufacturer’s protocol.

### X-ray crystallography

Crystallisation of bromodomain-peptide complexes was performed using a sitting-drop vapour-diffusion technique. Purified bromodomains (10–15 mg/mL) were combined with ∼1.5 molar equivalents of peptide and incubated on ice for at least 0.5 h. In situations where the bromodomain concentration was diluted below 5 mg/mL after addition of peptide, the bromodomain-peptide mixture was concentrated to bring the bromodomain concentration back to ∼10 mg/mL. Initial crystallisation trials were performed using commercial 96-well crystallisation screens. Bromodomain-peptide mixtures were dispensed into MRC two-drop chamber, 96-well crystallisation plates using a Mosquito crystallisation robot and each condition was screened at a 1:1 or 2:1 protein to precipitant ratio (maintaining a final drop volume of 300 nL). In certain cases, crystallisation was performed without pre-mixing of the bromodomain and peptides by separately dispensing 100 nL of protein and peptide directly into the crystallisation plate prior to mixing with the precipitant. Where required, initials hits were optimised by gradient refinement of the original condition, scaling up drop sizes, and microcrystal seeding. All experiments were performed at 18 °C. Protein crystals generally took days to weeks to appear. Crystals were frozen by plunge-freezing in liquid nitrogen following cryoprotection with 10% glycerol in the mother liquid from which the crystals were grown.

X-ray diffraction data were collected from frozen crystals at the Australian Synchrotron using the Macromolecular Crystallography MX1 (bending magnet) and MX2 beamlines (microfocus) at 100 K and a wavelength of 0.9537 Å^38, 39^. Data were integrated using XDS and were processed further using the CCP4i suite^40, 41^. AIMLESS was used for indexing, scaling, and merging of the data and the initial phases were calculated by the molecular replacement program PhaserMR using existing x-ray structures of bromodomains as the molecular replacement models (PDB IDs 4UYF for BRD2-BD1, 3ONI for BRD2-BD2, 3S91 for BRD3-BD1, 3S92 for BRD3-BD2, 4LYI for BRD4-BD1, and 5UVV for BRD4-BD2)^21, 23, 24, 42, 43^. Manual model building was performed using COOT and refinement was performed by iterative rounds of manual building in COOT followed by refinement using Phenix^42, 44^. The quality of the final model was validated with the wwPDB server and submitted to the PDB. Structure diagrams were generated using PyMol. Protein:peptide interfaces were analysed using PDBePISA. The data collection and refinement statistics for all structures described in this study are outlined in Supplementary Tables 2-16.

### NMR spectroscopy

NMR samples of BET BDs were prepared at ∼50-100 µM concentrations and were titrated with different molar equivalents of peptides. NMR spectra were acquired at 298 K using Bruker Avance III 600- or 800-MHz NMR spectrophotometers fitted with TCI probe heads and using standard pulse sequences from the Bruker library. TOPSPIN3 (Bruker) and NMRFAM-SPARKY were used for analysis of spectra^45^. Spectra were internally referenced to 10 μM 4,4-dimethyl-4-silapentane-1-sulfonic acid. Chemical shift perturbation experiments were performed by collecting ^15^N-HSQC spectra of ^15^N-labelled bromodomains before and after titration of unlabelled peptide into the samples. Interactions were assessed by monitoring the chemical shift perturbations induced by addition of the peptides to the bromodomains.

Chemical shift assignments were made for BRD3-BD1 using the standard triple resonance approach. Chemical Shift Perturbation (CSP) plots were generated using the following approach. A reference HSQC was recorded without the ligand present (ref HSQC), and then the ligand was added and a second HSQC (shift HSQC) recorded. The two HSQCs were peak picked using the APES algorithm in NMRFAM-Sparky to generate peak lists. The distance between each peak on the ref HSQC and all peaks on the shift HSQC was computed as follows:

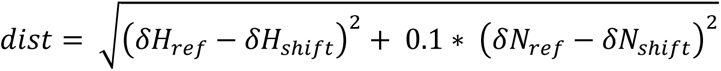

The minimum of the distances computed for each reference peak was taken as the CSP. Although this approach does not ensure every peak is correctly matched, it has the desirable property that the reported CSPs are guaranteed to be real and not an artefact of the algorithm, as the reported CSPs are a minimum of all possible CSP allocations.

The structure of **3.1B** was determined using NOESY, TOCSY and DQF-COSY spectra recorded at 5 °C. Spectra were analysed using CCPNMR Analysis and structures were calculated in CYANA3.98.5^46, 47^. Cyclization was introduced according to the protocol provided at the CYANA wiki (http://www.cyana.org/wiki/). The topology for the acetyllysine residues was created manually. For structures run with hydrogen bond restraints, the hydrogen bonds introduced were (a) Trp9-HN to Lys2 O and (b) Ile4 HN to AcK7 O. The restraints were added following the CYANA wiki recommendations. The final ensemble was the lowest-energy set of 20 models from 1000 calculated models.

### SEC-MALLS

Molecular weight calculations were carried out by subjecting samples (100 μL of 100-300 µM protein) to SEC (Superdex 75 10/300 Increase) on an Äkta system with inline multi-angle laser light (MALLS), UV absorbance, and refractive index (dRI) detectors. MALLS, UV, and dRI data were collected and analysed using the ASTRA software (Wyatt) and molecular weight determination was carried out according to the Debye-Zimm model. BSA (100 µL of 2 mg/mL) was run as a standard to calibrate align the MALLS, UV, and dRI signals. We estimate the uncertainty in the molecular weight determination from this system to be ±10%.

## Acknowledgements

JPM, LJW and RJP received funding from the National Health and Medical Research Council (APP 1161623). LJW received funding from the European Union’s Horizon 2020 research and innovation programme under the Marie Skłodowska-Curie grant agreement No 657292. This work was supported by Japan Agency for Medical Research and Development (AMED), Platform Project for Supporting Drug Discovery and Life Science Research (Basis for Supporting Innovative Drug Discovery and Life Science Research) under JP19am0101090 to HS. This work was also supported by the Francis Crick Institute which receives its core funding from Cancer Research UK (FC001748), the UK Medical Research Council (FC001748), and the Wellcome Trust (FC001748). This research was undertaken using MX1 and MX2 beamlines at the Australian Synchrotron, part of ANSTO, and made use of the Australian Cancer Research Foundation (ACRF) detector. We acknowledge the use of the Bosch Molecular Biology Facility at the University of Sydney for providing access to SPR infrastructure. We also thank the Crick Structural Biology Technology Platform for expert technical support and access to SPR infrastructure.

## Author Contributions

J.P.M., L.J.W., and T.P. conceived and J.P.M., L.J.W., T.P., H.S., R.J.P., J.M.G., and J.K.K.L supervised this study. J.K.K.L cloned constructs and K.P., J.L.W., and K.S.M expressed and purified proteins for this study. L.J.W. performed RaPID screens and L.J.W and D.T. synthesised peptides. L.J.W., K.P., and L.W. performed and analysed SPR experiments. J.M.M. contributed to binding data analysis. K.P., J.L.W., and K.S.M performed and analysed X-ray crystallography experiments. A.P.G.S. analysed x-ray data. K.P performed and analysed NMR and SEC-MALLS experiments. P.S. collected data for and solved NMR structures. J.P.M., L.J.W., T.P., and K.P prepared the manuscript with input from all authors.

