## Supplementary Figures for "Cyclic peptides can engage a single binding pocket through multiple, entirely divergent modes"

a

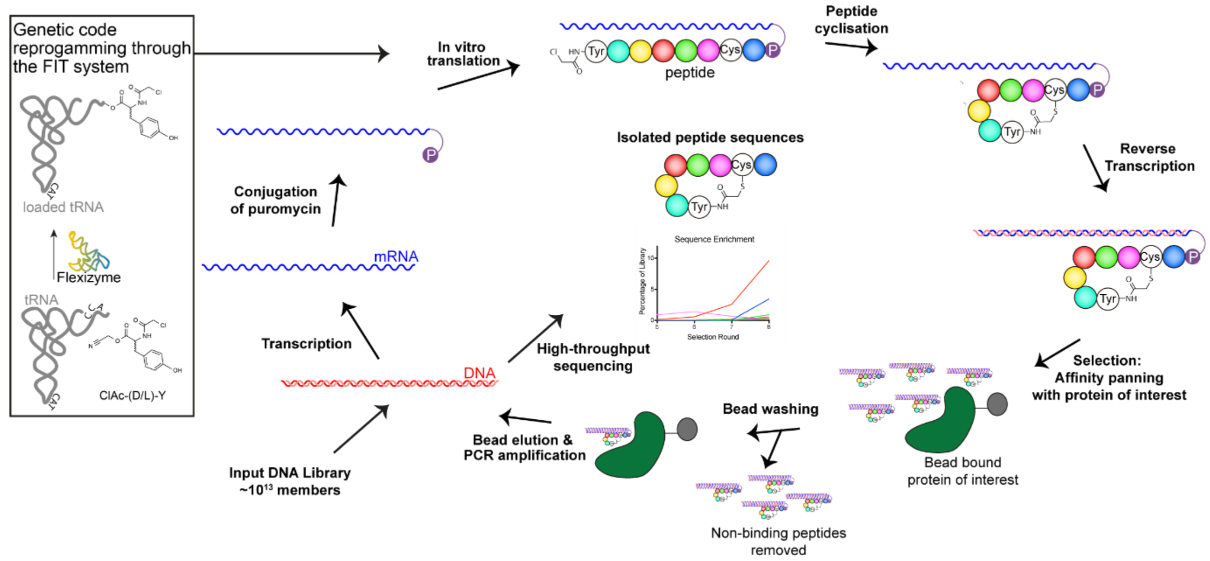


b

**m = 3-7**

**n = 4-7**

**ATG(NNK)mATG(NNK)nTGT**GGGTCTGGGTCTGGGTCTTAG

Peptide Sequence — Linker — Stop Codon

| 1st | 2nd U | 2nd C | 2nd A | 2nd G | 3rd |
|---|---|---|---|---|---|
| U | Phe | Ser | Tyr | Cys | U |
| | | | | | C |
| | | | | | A |
| | Leu | Ser | Stop | Trp | G |
| C | Leu | Pro | His | Arg | U |
| | | | | | C |
| | | | | | A |
| | Leu | Pro | Gln | Arg | G |
| A | Ile | Thr | Asn | Ser | U |
| | | | | | C |
| | | | | | A |
| | **KAc** | Thr | Lys | Arg | G |
| G | Val | Ala | Asp | Gly | U |
| | | | | | C |
| | | | | | A |
| | Val | Ala | Glu | Gly | G |

Initiator = ClAc-Trp

**Supplementary Figure 1.** Overview of the RaPID system. A. Schematic of the RaPID selection scheme. B. Codon assignment used in the RaPID selections.

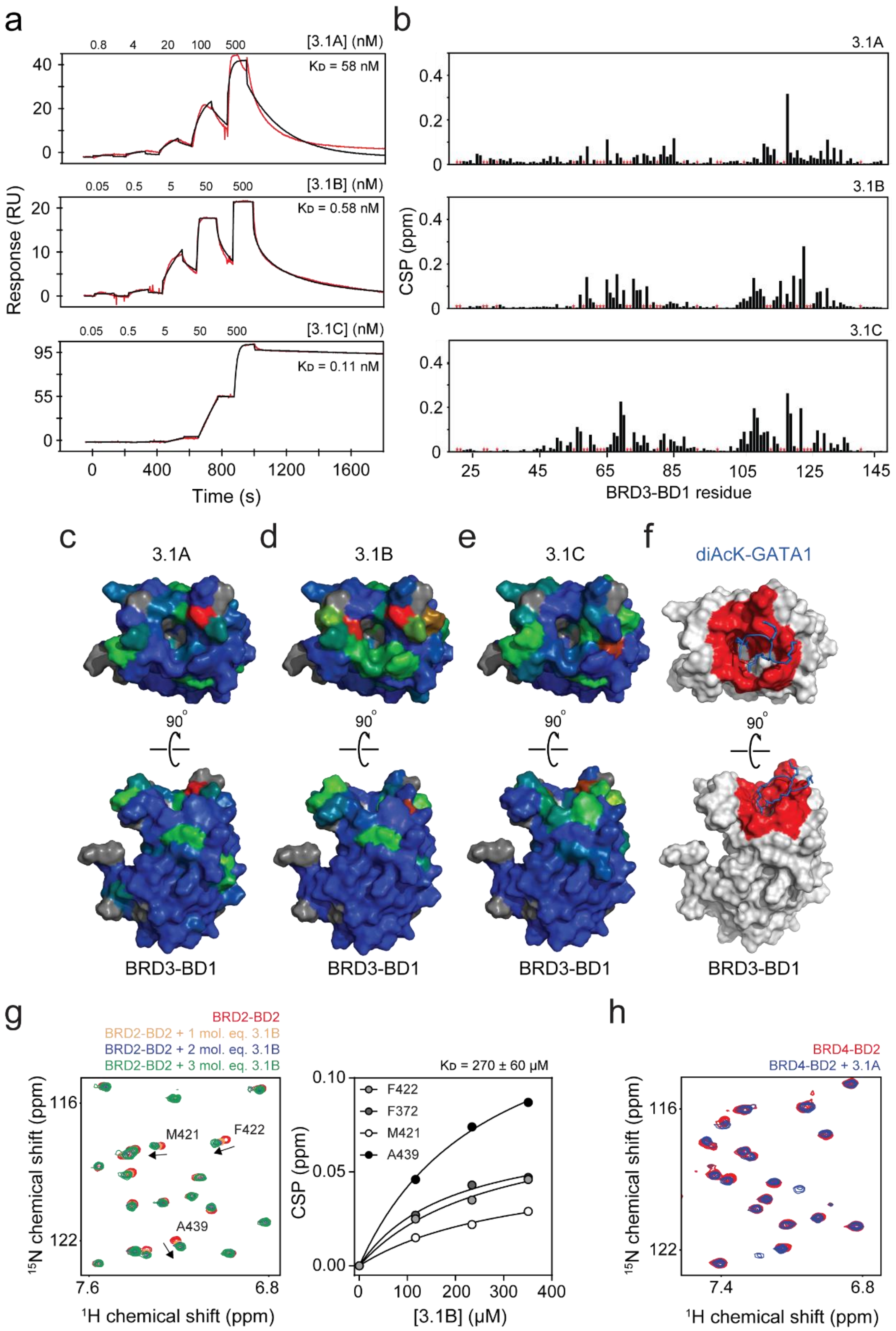


**Supplementary Figure 2. A. RaPID peptides bind BRD3-BD1 in the canonical AcK-binding pocket. a.** Representative SPR sensorgrams for BRD3-BD1 binding to **3.1A** (*upper panel*), **3.1B** (*middle panel*), and **3.1C** (*bottom panel*). **b.** Chemical shift perturbations in the $^{15}$N-HSQC of BRD3-BD1 upon formation of complexes with either **3.1A** (*upper panel*), or **3.1B** (*middle panel*), or **3.1C** (*lower panel*). The red stars indicate unassigned residues. **c.** CSPs observed upon formation of the **3.1A** complex mapped onto the structure of BRD3-BD1 (PDB: 3S91)[1]. The degrees of CSP are displayed through a blue-green-red gradient: the blue portions of the structure do not undergo any CSPs and the red sections represent the residues that display the greatest CSPs. Grey regions indicate unassigned residues. **d.** CSPs observed upon the **3.1B** complex mapped onto the structure of BRD3-BD1 (PDB: 3S91)[1]. The degrees of CSP are displayed through a blue-green-red gradient: the blue portions of the structure do not undergo any CSPs and the red sections represent the residues that display the greatest CSPs. Grey regions indicate unassigned residues. **e.** CSPs from the **3.1C** complex are mapped onto the structure of BRD3-BD1 (PDB: 3S91)[1]. The degrees of CSP are displayed through a blue-green-red gradient: the blue portions of the structure do not undergo any CSPs and the red sections represent the residues that display the greatest CSPs. Grey regions indicate unassigned residues. **f.** Structure of the BRD3-BD1 complex with a diacetylated-GATA1 peptide (2L5E, the peptide is shown in *blue* and the BD is shown in *grey*)[2], highlighting BD residues (*red*) that directly contact the peptide. **g.** Overlaid section of $^{15}$N-HSQC spectra of BRD2-BD2 alone (*red*) or in the presence of one (*yellow*), two (*blue*), or three (*green*) molar equivalents of **3.1B**. Signals that undergo the greatest CSPs in the are labelled in the protein alone spectrum. Binding curves were derived by tracking the combined $^{15}$N/$^{1}$H chemical shift perturbations of the signals undergoing the greatest CSPs. The data were fitted to a simple 1:1 Langmuir binding isotherm using Graphpad. **h.** Overlaid $^{15}$N-HSQC spectra of BRD4-BD2 alone (*red*) or in the presence of one (*blue*) molar equivalent of **3.1A**.

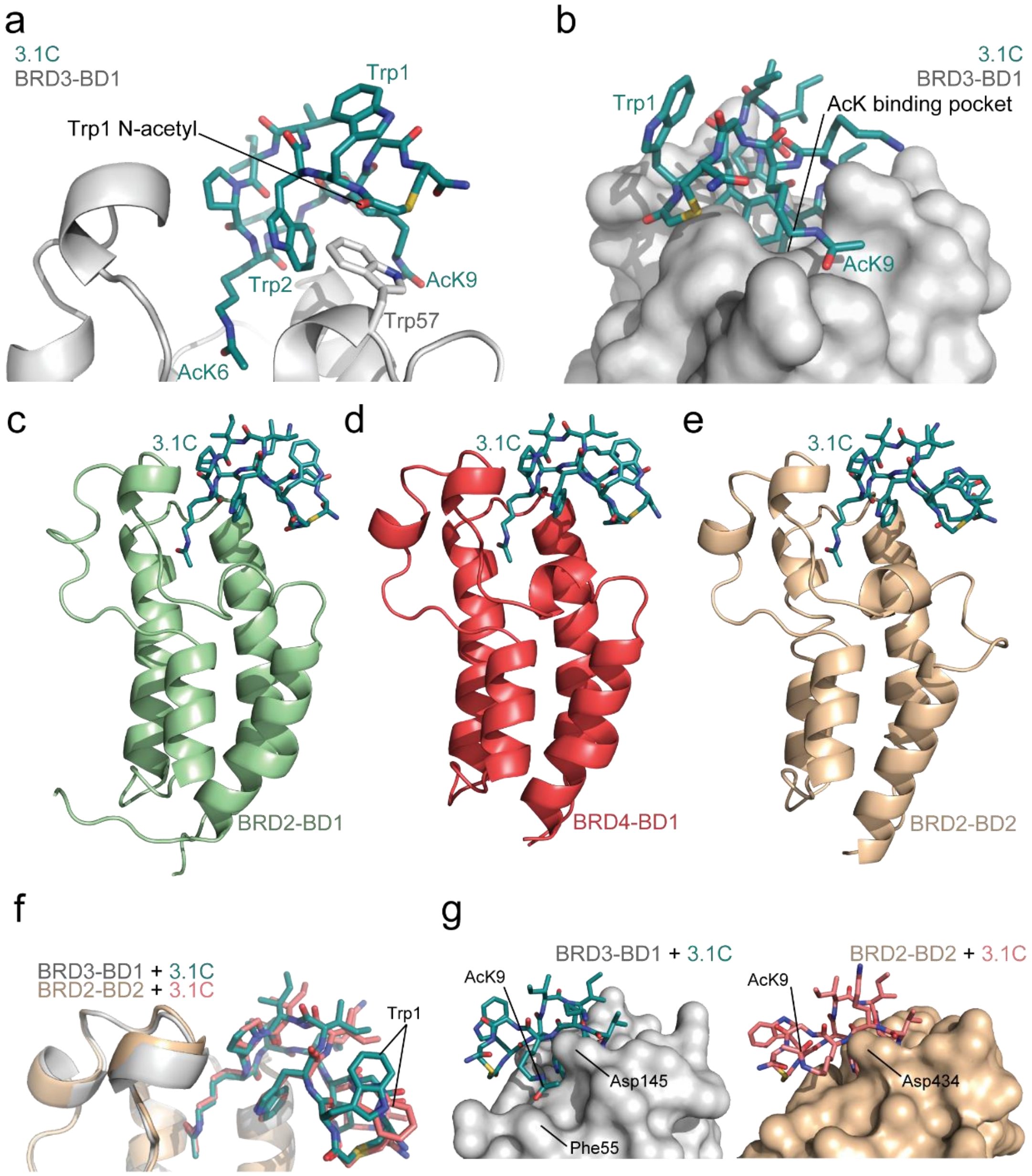


**Supplementary Figure 3. X-ray crystal structures of BRD2-BD1, BRD4-BD1 and BRD2-BD2 bound to 3.1C. a.**

**a.** The structure of BRD3-BD1 (*grey*) in complex with **3.1C** (*teal*) showing the sandwich of Trp57 from the BD between the Trp2 sidechain and the N-acetyl moiety of Trp1 from **3.1C**. **b.** Surface representation of BRD3-BD1 (*grey*) when in complex with **3.1C** (*teal*). AcK9 of **3.1C** interacts with a groove formed on the surface of the BD between the αZ and αD helices at a location distal to the AcK binding pocket. **c.** Ribbon representation of Ribbon representation of the X-ray crystal structure of BRD2-BD1 (*pale green*) bound to **3.1C** (*teal,* 2.3 Å resolution, PDB ID 6U61). **d.** Ribbon representation of the X-ray crystal structure of BRD4-BD1 (*coral red*) bound to **3.1C** (*teal,* 1.7 Å resolution, PDB IDs 6U6K). **e.** Ribbon representation of the X-ray crystal structure of BRD2-BD2 (*wheat*) bound to **3.1C** (*teal,* 1.5 Å resolution, PDB ID 6U71). **f.** Overlay of the BRD3-BD1:**3.1C** complex with the BRD2-BD2:**3.1C** complex. One of the major differences between the two peptide structures, the 180° flipping of the Trp1 sidechain, is indicated. **g.** Comparison of the orientation of AcK9 of **3.1C** and the surface representation the two BDs from the BRD3-BD1:**3.1C** and BRD2-BD2:**3.1C** complexes. BRD2-BD2 lacks the groove that AcK9 rests upon in the BRD3-BD1:**3.1C** structure leading to the residue being oriented upwards into the solvent in the BRD2-BD2:**3.1C** structure.

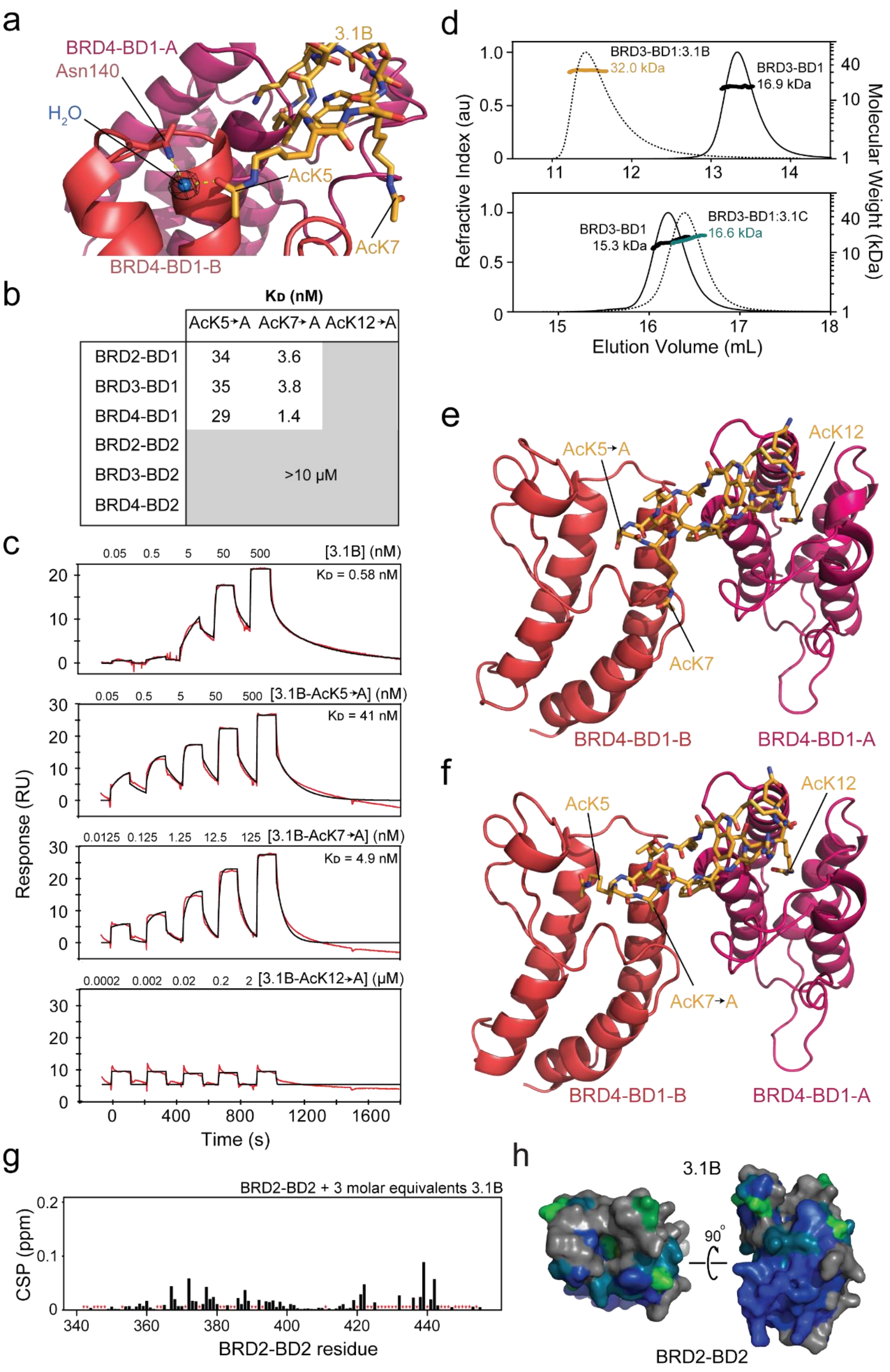


**Supplementary Figure 4. Analysis of BRD3/4-BD1:3.1B complexes. a.** A close up view of the water-mediated interaction between Asn140 of BRD4-BD1-B (*coral red*) and AcK5 of **3.1B** (*orange*). AcK5

enters the binding pocket at a distinct angle to the canonical AcK binding mode and forms a hydrogen bond with a water molecule (shown as a *blue sphere*) which in turn forms a hydrogen bond with Asn140. The electron density for the water is shown and hydrogen bonding is indicated by yellow dashed lines. **b.** Dissociation constants for the binding of each of the six BDs to **3.1B** AcK mutants. **c.** SPR data (*red*) for the binding of BRD3-BD1 to **3.1B** (*top*) and to AcK5→Ala, AcK7→Ala and AcK12→Ala mutants. Fits to a simple 1:1 binding model (*black*) are shown and derived $K_D$s are indicated on each plot. **d.** *Top:* Size-exclusion chromatogram for BRD3-BD1 alone (solid line) and in the presence of one molar equivalent of **3.1B** (dashed line). The calculated molecular weight trace from MALLS analysis is also shown. *Bottom:* Size-exclusion chromatogram for BRD3-BD1 alone (solid line) and in the presence of one molar equivalent of **3.1C** (dashed line). The calculated molecular weight trace from MALLS analysis is also shown. BRD3-BD1 alone elutes as a single peak with the expected mass of 17 kDa. Addition of one molar equivalent of **3.1B** (Mw =1.9 kDa) shifts the peak to an earlier elution time and yields a mass of 32 kDa, in close agreement with the expected mass of a 2:1 BD-**3.1B** complex. **e.** Ribbon diagram of the BRD4-BD1:**3.1B**_AcK5→Ala X-ray crystal structure (2.3 Å resolution, PDB ID 6U72). The peptide is shown in *orange* and the two BDs bound to the peptide are shown in pink (BRD4-BD1-A) and *coral red* (BRD4-BD1-B). The AcK5→Ala replacement and remaining AcKs are indicated. **f.** Ribbon diagram of the BRD4-BD1:**3.1B**_AcK7→Ala X-ray crystal structure (2.6 Å resolution, PDB ID 6U8G). The peptide is shown in *orange* and the two BDs bound to the peptide are shown in pink (BRD4-BD1-A) and *coral red* (BRD4-BD1-B). The AcK7→Ala replacement and remaining AcKs are indicated. **g.** Chemical shift perturbations in the $^{15}$N-HSQC of BRD2-BD2 upon formation of complexes following addition of three molar equivalents of **3.1B**. The red stars indicate unassigned residues. **h.** CSPs observed upon formation of the **3.1B** complex mapped onto the structure of BRD2-BD2 (PDB: 3ONI)[1]. The degrees of CSP are displayed through a blue-green-red gradient: the blue portions of the structure do not undergo any CSPs and the red sections represent the residues that display the greatest CSPs. Grey regions indicate unassigned residues.

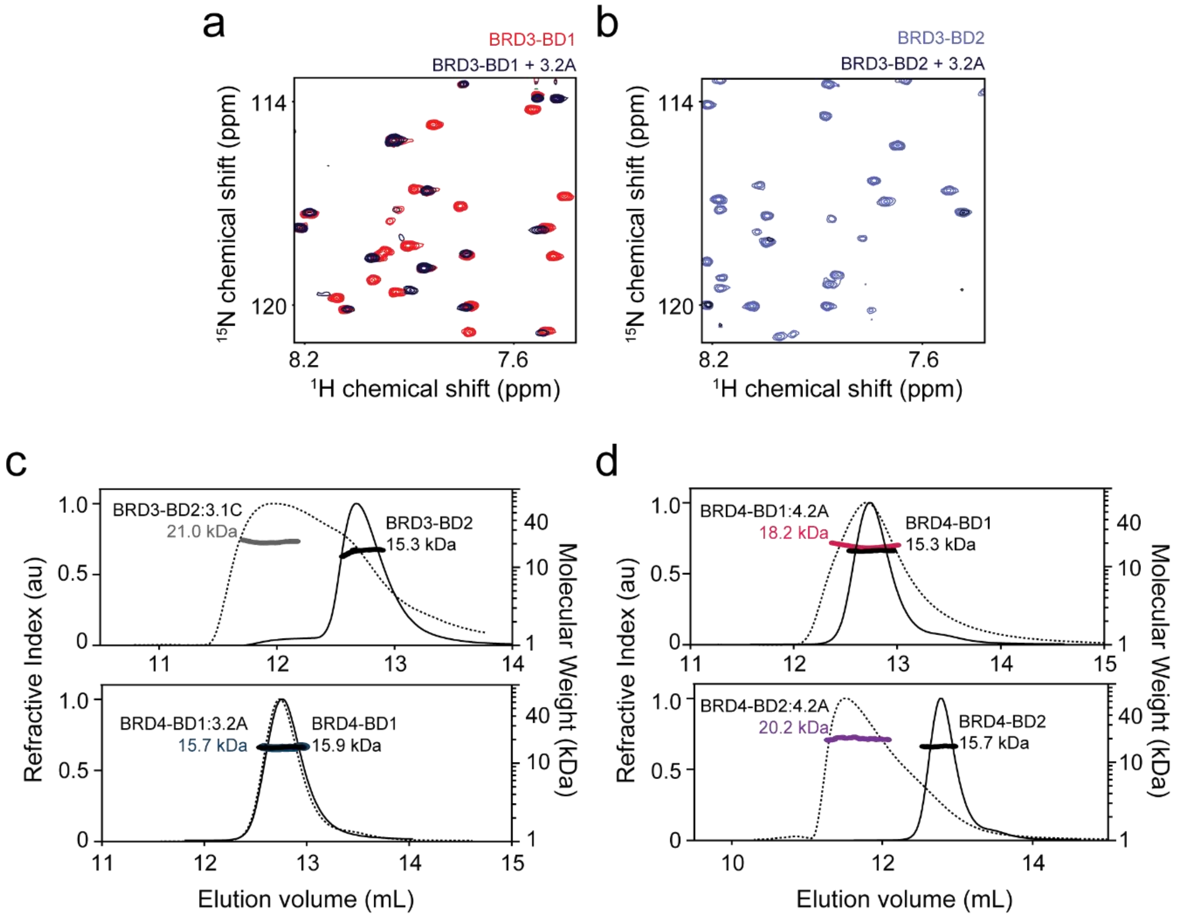


**Supplementary Figure 5. $^{15}$N-HSQC analysis of interactions with 3.2A. a.** Overlaid $^{15}$N-HSQC spectra of BRD3-BD1 alone (*red*) or in the presence of 1.5 molar equivalents of **3.2A** (*dark purple*). **b.** Overlaid $^{15}$N-HSQC spectra of BRD3-BD2 alone (*light purple*) or in the presence of 0.5 molar equivalents of **3.2A**. There is a mass disappearance of signals upon addition of **3.2A**. **c.** *Top:* Size-exclusion chromatogram for BRD3-BD2 alone (solid line) and in the presence of 0.5 molar equivalents of **3.2C** (dashed line). The calculated molecular weight from MALLS analysis is also shown. Addition of 0.5 molar equivalents of **3.2C** leads to a shift from a peak that yields a mass of 15.3 kDa (BRD3-BD2 alone) to species with an earlier elution time that yields a molecular weight of 21.0 kDa, suggesting the formation of a weak 2:1 BD:**3.2C** complex. *Bottom:* Size-exclusion chromatogram for BRD4-BD1 alone (solid line) and in the presence of 0.5 molar equivalents of **3.2A** (dashed line). The calculated molecular weight from MALLS analysis is also shown. **d.** *Top:* Size-exclusion chromatogram for BRD4-BD1 alone (solid line) and in the presence of 0.5 molar equivalent of **4.2A** (dashed line). The calculated molecular weight trace from MALLS analysis is also shown. *Bottom:* Size-exclusion chromatogram for BRD4-BD2 alone (solid line) and in the presence of one molar equivalent of **3.1C** (dashed line). The calculated molecular weight trace from MALLS analysis is also shown. BRD4-BD1 alone elutes as a single peak with the expected mass of 15.3 kDa. Addition of 0.5 molar equivalent of **4.2A** causes peak broadening and yields a mass of 18.3 kDa, consistent with a 1:1 BD:**4.2A** complex. Addition of 0.5 molar equivalents of **4.2A** to BRD4-BD2 leads to a shift from a peak that yields a mass of 15.7 kDa (BRD4-BD2 alone) to a species with an earlier elution time that yields a molecular weight of 20.2, indicating the formation of a weak 2:1 BD:**4.2A** complex.

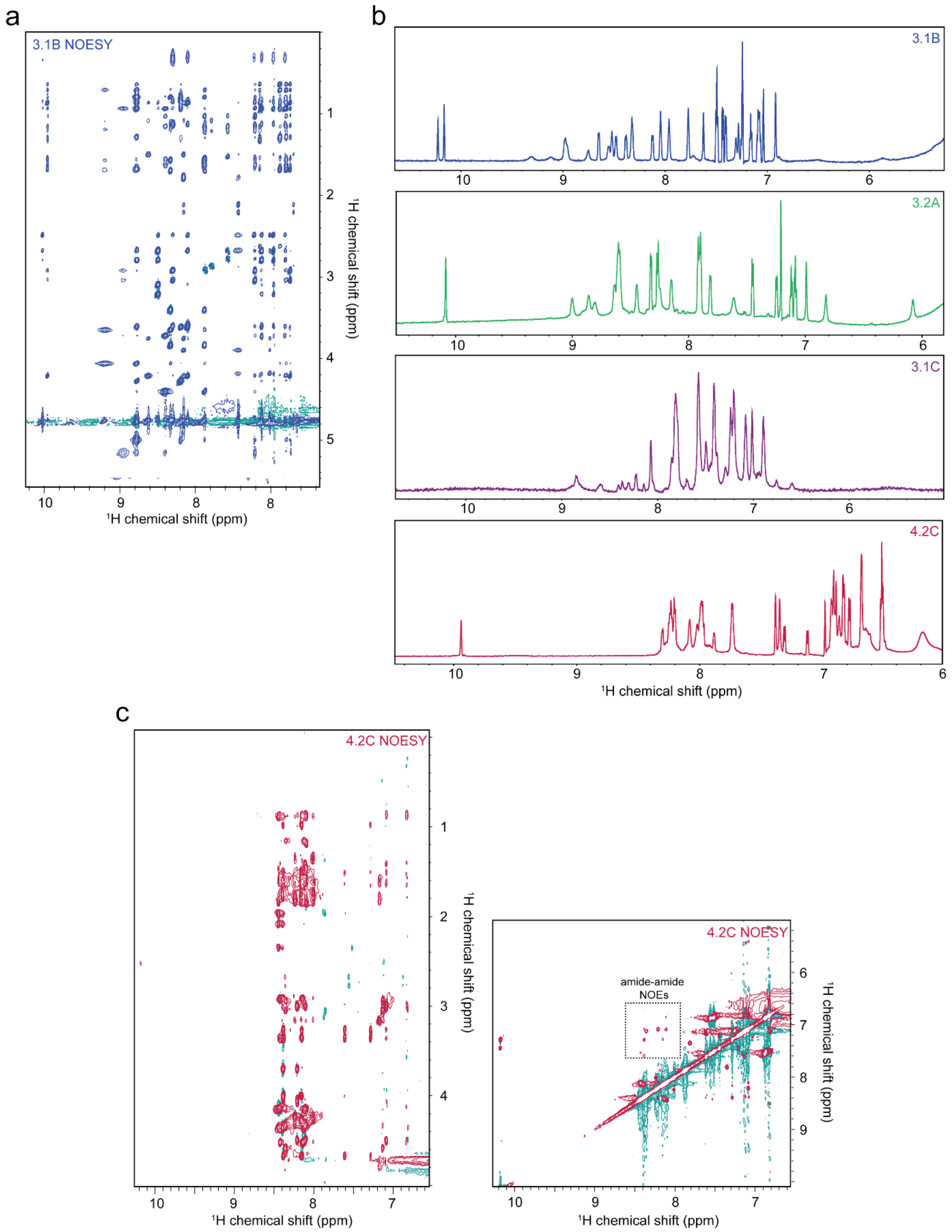


**Supplementary Figure 6. Two-dimensional NOESY and one-dimensional $^{1}$H-NMR spectra of selected peptides**. **a.** Section of a two-dimensional NOESY spectra of **3.1B**. **b.** One-dimensional $^{1}$H-NMR spectra of **3.1B**, **3.2A**, **3.1C**, and **4.2C**. **c.** Sections of two-dimensional NOESY spectra of **4.2C**. All spectra were collected at 25 °C on an 800-MHz spectrometer.

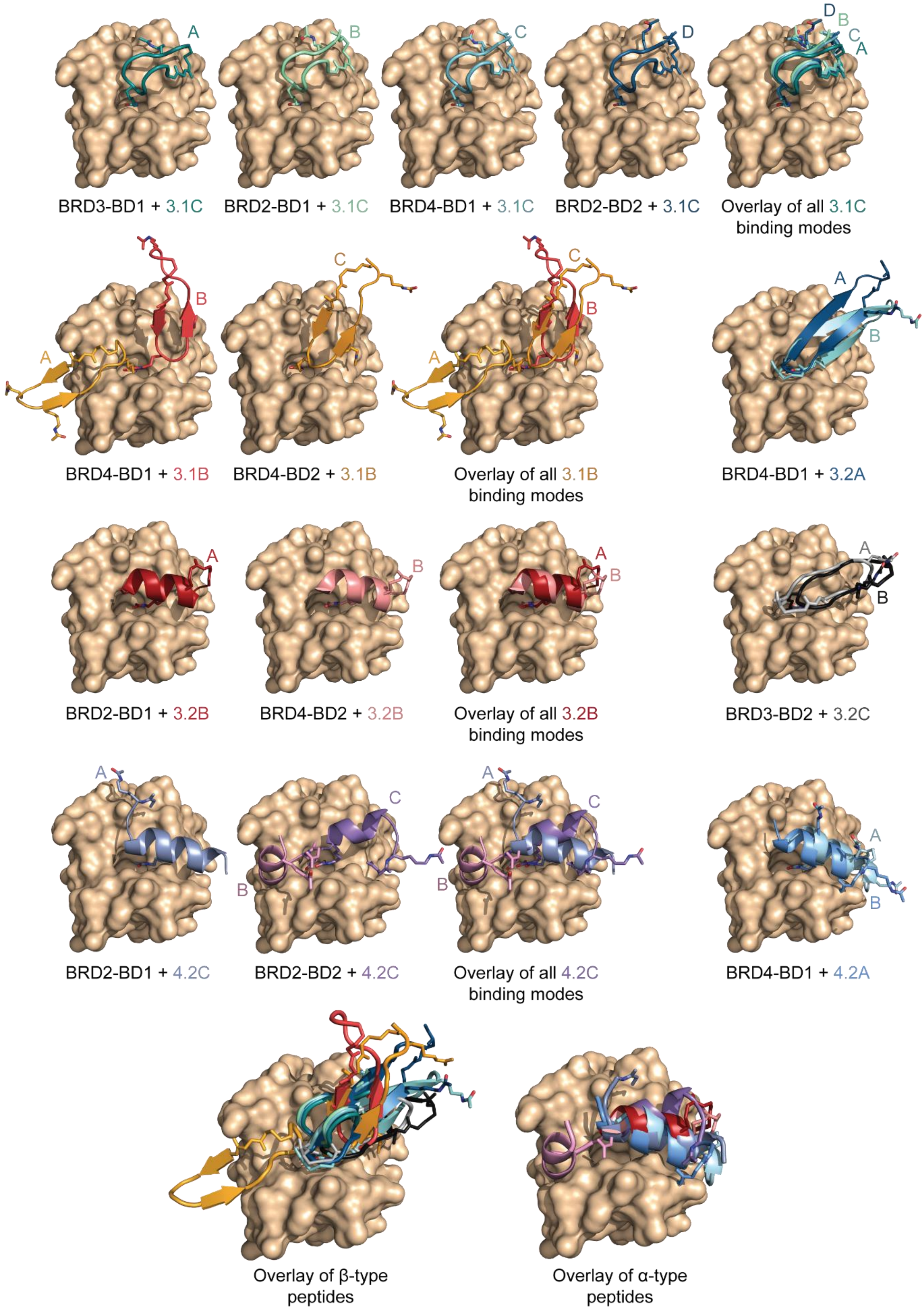


**Supplementary Figure 7. RaPID cyclic peptide structure summary**

3.1A

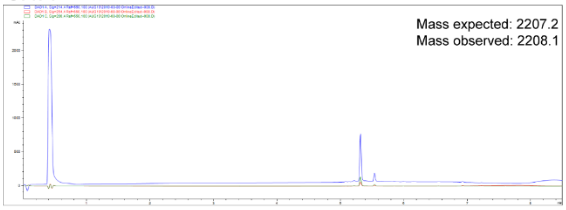


3.2A

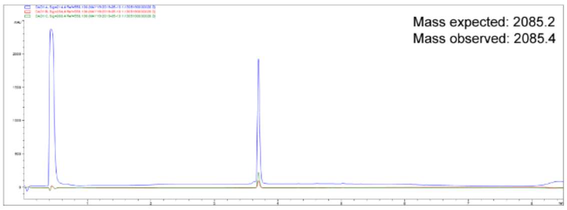


3.1B

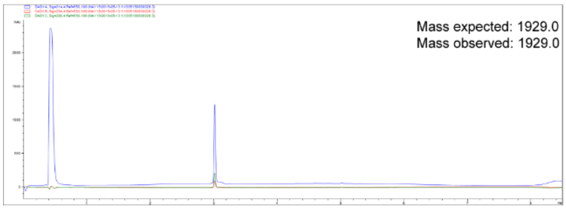


3.2B

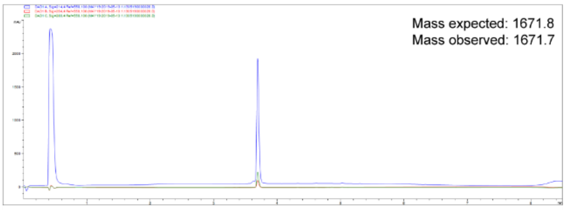


3.2C

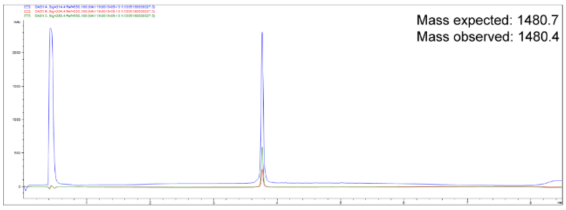


3.2C

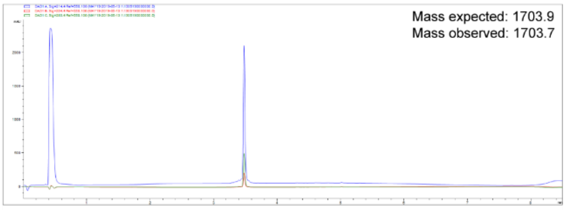


4.2A

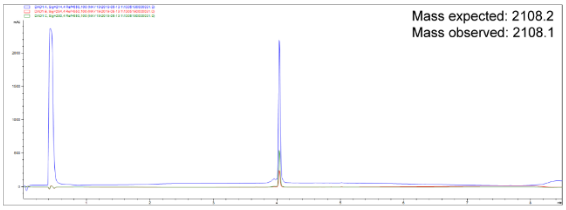


3.1B AcK5Ala

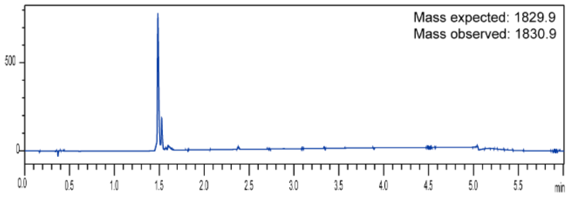


4.2B

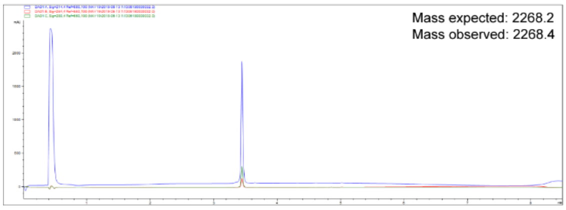


3.1B AcK7Ala

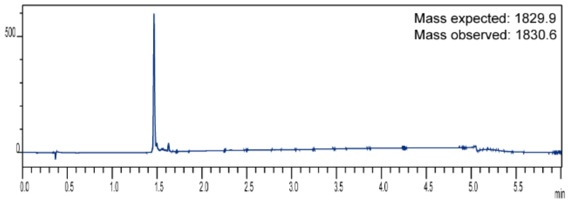


4.2C

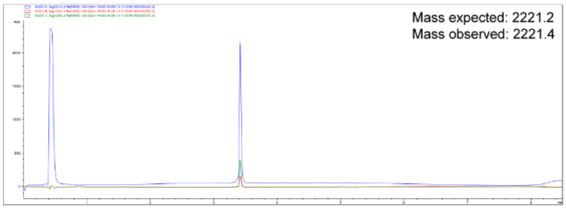


**Supplementary Figure 8. HPLC traces of all RaPID peptides**
