## Supplementary Tables 2-10 for "Cyclic peptides can engage a single binding pocket through multiple, entirely divergent modes"

**Supplementary Table 2.**

Data collection and refinement statistics for the crystal structures of the BRD3-BD1:**3.1C** (PDB ID 6U4A) and BRD2-BD1:**3.1C** (PDB ID 6U61) complexes solved by molecular replacement.

| | BRD3-BD1:**3.1C** | BRD2-BD1:**3.1C** |
|---|---|---|
| **Data collection** | | |
| Space group | P 21 21 21 | C 2 2 21 |
| Cell dimensions | | |
| *a*, *b*, *c* (Å) | 48.83, 64.42, 87.65 | 69.54, 86.75, 115.48 |
| α, β, γ (°) | 90.00, 90.00, 90.00 | 90.00, 90.00, 90.00 |
| Resolution (Å) | 1.88 (1.88-1.92) | 2.29 (2.29-2.37) |
| $R_{merge}$ | 0.075 (0.515) | 0.113 (0.775) |
| $I / \sigma I$ | 16.4 (3.8) | 13.9 (4.3) |
| Completeness (%) | 99.9 (0.883) | 99.6 (96.4) |
| Redundancy | 7.4 (7.2) | 13.2 (12.7) |
| **Refinement** | | |
| Resolution (Å) | 1.88 | 2.29 |
| No. reflections | 23024 | 16071 |
| $R_{work}$ / $R_{free}$ | 0.1637/0.1881 | 0.222/0.2634 |
| No. atoms | | |
| Protein | 2268 | 2165 |
| Ligand/ion | 36 | 11 |
| Water | 206 | 77 |
| *B*-factors | 24.0 | 43.0 |
| R.m.s. deviations | | |
| Bond lengths (Å) | 0.004 | 0.002 |
| Bond angles (°) | 0.674 | 0.519 |

*Values in parentheses are for highest-resolution shell.
All were data were collected from a single crystal.

**Supplementary Table 3.**

Data collection and refinement statistics for the crystal structures of the BRD4-BD1:**3.1C** (PDB ID 6U6K) and BRD2-BD2:**3.1C** (PDB ID 6U71) complexes solved by molecular replacement.

| | BRD4-BD1:**3.1C** | BRD2-BD2:**3.1C** |
|---|---|---|
| **Data collection** | | |
| Space group | C 1 2 1 | P 2 21 21 |
| Cell dimensions | | |
| *a*, *b*, *c* (Å) | 121.96, 41.90, 29.54 | 31.82, 51.81, 71.79 |
| α, β, γ (°) | 90.00, 94.20, 90.00 | 90.00, 90.00, 90.00 |
| Resolution (Å) | 1.70 (1.70-1.73) | 1.47 (1.47-1.50) |
| $R_{merge}$ | 0.061 (0.611) | 0.062 (0.547) |
| $I / \sigma I$ | 9.0 (1.8) | 11.8 (2.7) |
| Completeness (%) | 99.7 (99.7) | 99.7 (99.3) |
| Redundancy | 3.3 (3.2) | 4.4 (4.4) |
| **Refinement** | | |
| Resolution (Å) | 1.70 | 1.47 |
| No. reflections | 16475 | 20844 |
| $R_{work}$ / $R_{free}$ | 0.1799/0.2177 | 0.1600/0.1991 |
| No. atoms | | |
| Protein | 1152 | 1150 |
| Ligand/ion | 4 | 4 |
| Water | 110 | 116 |
| *B*-factors | 27.0 | 20 |
| R.m.s. deviations | | |
| Bond lengths (Å) | 0.007 | 0.003 |
| Bond angles (°) | 0.934 | 0.615 |

*Values in parentheses are for highest-resolution shell.
All were data were collected from a single crystal.

**Supplementary Table 4.**

Data collection and refinement statistics for the crystal structures of the BRD4-BD1:**3.1B** (PDB ID 6U74) and BRD4-BD1:**3.1B**_AcK5→Ala (PDB ID 6U72) complexes solved by molecular replacement.

| | BRD4-BD1:**3.1B** | BRD4-BD1:**3.1B**_AcK5→Ala |
|---|---|---|
| **Data collection** | | |
| Space group | P 1 21 1 | P 1 21 1 |
| Cell dimensions | | |
| *a*, *b*, *c* (Å) | 58.68, 49.23, 115.94 | 58.21, 49.05, 59.15 |
| α, β, γ (°) | 90.00, 102.58, 90.00 | 90.00, 103.87, 90.00 |
| Resolution (Å) | 1.85 (1.85-1.89) | 2.30 (2.30-2.38) |
| $R_{merge}$ | 0.317 (2.03) | 0.145 (0.638) |
| $I / \sigma I$ | 4.5 (1.0) | 7.8 (2.7) |
| Completeness (%) | 100.0 (100.0) | 99.9 (99.9) |
| Redundancy | 7.4 (7.4) | 6.7 (7.0) |
| **Refinement** | | |
| Resolution (Å) | 1.85 | 2.30 |
| No. reflections | 54657 | 14555 |
| $R_{work}$ / $R_{free}$ | 0.2841/0.3234 | 0.2093/0.2661 |
| No. atoms | | |
| Protein | 4161 | 2074 |
| Ligand/ion | 8 | 4 |
| Water | 457 | 63 |
| *B*-factors | 27.0 | 59.0 |
| R.m.s. deviations | | |
| Bond lengths (Å) | 0.002 | 0.003 |
| Bond angles (°) | 0.570 | 0.533 |

*Values in parentheses are for highest-resolution shell.
All were data were collected from a single crystal.

**Supplementary Table 5.**

Data collection and refinement statistics for the crystal structures of the BRD4-BD1:**3.1B**_AcK7→Ala (PDB ID 6U8G) and BRD4-BD2:**3.1B** (PDB ID 6U6L) complexes solved by molecular replacement.

| | BRD4-BD1:**3.1B**_AcK7→Ala | BRD4-BD2:**3.1B** |
|---|---|---|
| **Data collection** | | |
| Space group | P 1 21 1 | P 2 21 21 |
| Cell dimensions | | |
| *a*, *b*, *c* (Å) | 58.71, 48.98, 114.15 | 32.40, 56.51, 73.34 |
| α, β, γ (°) | 90.00, 101.66, 90.00 | 90.00, 90.00, 90.00 |
| Resolution (Å) | 2.6 (2.6-2.72) | 2.6 (2.6-2.72) |
| $R_{merge}$ | 0.237 (1.021) | 0.103 (0.636) |
| $I / \sigma I$ | 6.1 (2.6) | 12.6 (2.8) |
| Completeness (%) | 99.8 (100.0) | 97.3 (98.6) |
| Redundancy | 5.7 (5.6) | 4.7 (4.7) |
| **Refinement** | | |
| Resolution (Å) | 2.60 | 2.60 |
| No. reflections | 19831 | 4472 |
| $R_{work}$ / $R_{free}$ | 0.2416/0.2965 | 0.2160/0.2425 |
| No. atoms | | |
| Protein | 4169 | 1043 |
| Ligand/ion | 8 | 10 |
| Water | 45 | 16 |
| *B*-factors | 47.0 | 42.0 |
| R.m.s. deviations | | |
| Bond lengths (Å) | 0.001 | 0.002 |
| Bond angles (°) | 0.462 | 0.525 |

*Values in parentheses are for highest-resolution shell.
All were data were collected from a single crystal.

**Supplementary Table 6.**

Data collection and refinement statistics for the crystal structures of the BRD3-BD2:**3.2C** (PDB ID 6ULP) and BRD4-BD1:**3.2A** (PDB ID 6U8M) complexes solved by molecular replacement.

| | BRD3-BD2:**3.2C** | BRD4-BD1:**3.2A** |
|---|---|---|
| **Data collection** | | |
| Space group | I 4 | P 21 21 21 |
| Cell dimensions | | |
| *a*, *b*, *c* (Å) | 97.57, 97.57, 77.75 | 61.83, 72.62, 86.06 |
| α, β, γ (°) | 90.00, 90.00, 90.00 | 90.00, 90.00, 90.00 |
| Resolution (Å) | 2.80 (2.80-2.96) | 1.95 (1.95-2.00) |
| $R_{merge}$ | 0.049 (0.832) | 0.061 (0.432) |
| $I / \sigma I$ | 29.9 (3.4) | 22.1 (4.8) |
| Completeness (%) | 99.8 (98.8) | 100.0 (100.0) |
| Redundancy | 14.0 (14.2) | 13.0 (13.2) |
| **Refinement** | | |
| Resolution (Å) | 2.80 | 1.95 |
| No. reflections | 9017 | 28898 |
| $R_{work}$ / $R_{free}$ | 0.2102/0.2477 | 0.1700/0.1999 |
| No. atoms | | |
| Protein | 1932 | 2117 |
| Ligand/ion | 4 | 4 |
| Water | 0 | 214 |
| *B*-factors | 110.0 | 41.0 |
| R.m.s. deviations | | |
| Bond lengths (Å) | 0.002 | 0.008 |
| Bond angles (°) | 0.559 | 0.912 |

*Values in parentheses are for highest-resolution shell.
All were data were collected from a single crystal.

**Supplementary Table 7.**

Data collection and refinement statistics for the crystal structures of the BRD2-BD1:**3.2B** (PDB ID 6U8H) and BRD4-BD2:**3.2B** (PDB ID 6U8I) complexes solved by molecular replacement.

| | BRD2-BD1:**3.2B** | BRD4-BD2:**3.1B** |
|---|---|---|
| **Data collection** | | |
| Space group | P 41 21 2 | P 4 |
| Cell dimensions | | |
| *a*, *b*, *c* (Å) | 89.59, 89.59, 71.59 | 69.78, 69.78, 32.15 |
| α, β, γ (°) | 90.00, 90.00, 90.00 | 90.00, 90.00, 90.00 |
| Resolution (Å) | 2.07 (2.07-2.13) | 2.50 (2.50-2.70) |
| $R_{merge}$ | 0.062 (0.322) | 0.165 (0.593) |
| $I / \sigma I$ | 25.0 (7.9) | 6.4 (2.5) |
| Completeness (%) | 99.7 (97.0) | 100.00 (100.0) |
| Redundancy | 12.2 (12.0) | 4.9 (4.9) |
| **Refinement** | | |
| Resolution (Å) | 2.07 | 2.50 |
| No. reflections | 18148 | 5552 |
| $R_{work}$ / $R_{free}$ | 0.1819/0.2232 | 0.1892/0.2395 |
| No. atoms | | |
| Protein | 1169 | 1004 |
| Ligand/ion | 65 | 4 |
| Water | 129 | 59 |
| *B*-factors | 34.0 | 28.0 |
| R.m.s. deviations | | |
| Bond lengths (Å) | 0.002 | 0.002 |
| Bond angles (°) | 0.465 | 0.439 |

*Values in parentheses are for highest-resolution shell.
All were data were collected from a single crystal.

**Supplementary Table 8.**

Data collection and refinement statistics for the crystal structures of the BRD4-BD1:**4.2A** (PDB ID 6ULV) complex solved by molecular replacement.

| | BRD4-BD1:**4.2A** |
|---|---|
| **Data collection** | |
| Space group | P 65 2 2 |
| Cell dimensions | |
| *a*, *b*, *c* (Å) | 112.29, 112.29, 235.58 |
| α, β, γ (°) | 90.00, 90.00, 120.00 |
| Resolution (Å) | 2.20 (2.20-2.27) |
| $R_{merge}$ | 0.474 (7.386) |
| $I / \sigma I$ | 9.2 (1.4) |
| Completeness (%) | 100.0 (100.0) |
| Redundancy | 33.3 (34.0) |
| **Refinement** | |
| Resolution (Å) | 2.20 |
| No. reflections | 45317 |
| $R_{work}$ / $R_{free}$ | 0.1984 (0.2377) |
| No. atoms | |
| Protein | 3957 |
| Ligand/ion | 46 |
| Water | 240 |
| *B*-factors | 45.0 |
| R.m.s. deviations | |
| Bond lengths (Å) | 0.004 |
| Bond angles (°) | 0.682 |

*Values in parentheses are for highest-resolution shell.
All were data were collected from a single crystal.

**Supplementary Table 9.**

Data collection and refinement statistics for the crystal structures of the BRD2-BD1:**4.2C** (PDB ID 6ULT) and BRD2-BD2:**4.2C** (PDB ID 6ULQ) complexes solved by molecular replacement.

| | BRD2-BD1:**4.2C** | BRD2-BD2:**4.2C** |
|---|---|---|
| **Data collection** | | |
| Space group | I 1 2 1 | P 1 21 1 |
| Cell dimensions | | |
| *a*, *b*, *c* (Å) | 80.48, 76.14, 109.66 | 77.08, 63.63, 120.34 |
| α, β, γ (°) | 90.00, 109.28, 90.00 | 90.00, 108.39, 90.00 |
| Resolution (Å) | 2.70 (2.70-2.83) | 2.80 (2.80-2.95) |
| $R_{merge}$ | 0.277 (1.454) | 0.165 (0.686) |
| $I / \sigma I$ | 9.5 (2.9) | 6.4 (1.8) |
| Completeness (%) | 99.6 (97.6) | 98.8 (94.2) |
| Redundancy | 6.7 (6.9) | 3.7 (3.8) |
| **Refinement** | | |
| Resolution (Å) | 2.70 | 2.80 |
| No. reflections | 17421 | 27196 |
| $R_{work}$ / $R_{free}$ | 0.2215/0.2516 | 0.2814/0.3379 |
| No. atoms | | |
| Protein | 3550 | 7462 |
| Ligand/ion | 9 | 12 |
| Water | 36 | 81 |
| *B*-factors | 34.0 | 55.0 |
| R.m.s. deviations | | |
| Bond lengths (Å) | 0.002 | 0.003 |
| Bond angles (°) | 0.571 | 0.690 |

*Values in parentheses are for highest-resolution shell.
All were data were collected from a single crystal.

**Supplementary Table 10.**

NMR and refinement statistics for the solution structure of cyclic peptide **3.1B** (PDB ID: 6UXS).

| | **Protein** |
|---|---|
| **NMR distance and dihedral constraints** | |
| Distance constraints | |
| Total NOE | 261 |
| Intra-residue | 91 |
| Inter-residue | 170 |
| Sequential ($\|i-j\| = 1$) | 66 |
| Medium-range ($\|i-j\| < 4$) | 28 |
| Long-range ($\|i-j\| > 5$) | 76 |
| Intermolecular | 0 |
| Hydrogen bonds | 0 |
| Total dihedral angle restraints | |
| $\phi$ | 0 |
| $\psi$ | 0 |
| **Structure statistics** | |
| Violations (mean and s.d.) | |
| Distance constraints (Å) | 0 |
| Dihedral angle constraints (º) | N/A |
| Max. dihedral angle violation (º) | N/A |
| Max. distance constraint violation (Å) | 0 |
| Deviations from idealized geometry | |
| Bond lengths (Å) | 0.001 |
| Bond angles (º) | 0.2 |
| Impropers (º) | N/A |
| Average pairwise r.m.s. deviation** (Å) | |
| Heavy | 0.75 |
| Backbone | 0.02 |

*Pairwise r.m.s. deviation was calculated among 20 refined structures.
